# Developmental patterning of adipose tissue by *abd-A* and *Abd-B* homeotic genes in *Drosophila melanogaster*

**DOI:** 10.64898/2026.05.22.727268

**Authors:** Rajitha-Udakara-Sampath Hemba-Waduge, Mengmeng Liu, Xiao Li, Elisabeth A. Budslick, Sarah E. Bondos, Keith A. Maggert, Jun-Yuan Ji

## Abstract

The Bithorax Complex (BX-C) homeobox proteins specify segmental identities along the anterior-posterior axis during *Drosophila* embryogenesis. Differential expression of the *BX-C* genes *abd-A* and *Abd-B* distinguishes abdominal from thoracic adipocytes, yet the mechanism regulating this heterogeneity remains poorly understood. Here, we identify cis-regulatory elements (CREs) and transcription factors that direct abdominal-specific expression of *abd-A* and *Abd-B* in the larval fat body. Fine-mapping analyses identified a 627-bp CRE within the *Abd-B* locus and a ∼6-kb CRE within the *abd-A* locus sufficient to drive heterogeneous expression. Yeast one-hybrid screening combined with functional analyses identified Lola, Lolal, and Combgap as key repressors of *Abd-B*, whereas Piragua (Prg) and Seven up (Svp) function as transcriptional activators, indicating that adipocyte heterogeneity in postembryonic adipose tissue is actively regulated. In turn, *lola* and *prg* are repressed by Abd-B, whereas *lolal* and *svp* are activated, forming a feedback circuit further modulated by Wnt signaling, which promotes *lola* and *lolal* expression while repressing *svp*. CUT&RUN analyses suggest that these interactions are direct, with dTCF/Pan and Abd-B occupancy detected at target loci. Together, our findings define a transcriptional circuit that regulates *Abd-B* gene transcription to pattern adipose tissue and may establish the developmental basis of fat depot specialization.

**One-sentence summary:** Transcriptional regulation of *abd-A* & *Abd-B* in adipocytes

## Introduction

Pioneering studies using *Drosophila melanogaster* have elucidated fundamental mechanisms governing body plan formation in metazoans. These processes rely on precisely coordinated spatial and temporal regulation of maternal and zygotic gene expression following fertilization, culminating in the formation of a multicellular blastoderm (Saunders 2021; Martin 2020; Martin et al. 2009). Maternal effect genes, including *bicoid* and *nanos*, establish the anterior-posterior axis of the early embryo (Kimelman and Martin 2012; Saunders 2021), whereas segmentation genes, such as gap genes and pair-rule genes, further subdivide the embryo into discrete segments (Jaeger 2011; Diaz-Cuadros et al. 2021). These early patterning cues converge on the regulation of downstream target genes, most notably the *homeobox* (*Hox*) genes. Precise control of *Hox* gene expression is essential for proper axial patterning and the formation of segment-specific structures (Soshnikova and Duboule 2009; Vasanthi and Mishra 2008). During postembryonic development, Polycomb group (PcG) and Trithorax group (TrxG) proteins preserve *Hox* gene expression states via epigenetic modification of histone tails (Schuettengruber et al. 2017; Kassis et al. 2017). However, establishing a permissive chromatin landscape alone is insufficient to generate the precise spatial and temporal transcriptional patterns *in vivo*. Accurate expression requires the concerted action of sequence-specific transcription factors, co-activators, and co-repressors, which engage cis-regulatory elements (CREs), recruit the transcriptional machinery, and integrate contextual signaling cues.

In *Drosophila*, *Hox* genes are organized into two clusters on the third chromosome: the Antennapedia Complex (ANT-C) and the Bithorax Complex (BX-C). The ANT-C comprises five *Hox* genes: *labial* (*lab*), *proboscipedia* (*pb*), *Deformed* (*Dfd*), *Sex combs reduced* (*Scr*), and *Antennapedia* (*Antp*), which specify the identities of the anterior segments (Struhl 1981). The *BX-C* contains three *Hox* genes: *Ultrabithorax* (*Ubx*), *abdominal-A* (*abd-A*), and *Abdominal-B* (*Abd-B*), which define posterior segment identities (Lewis 1978). *Ubx* specifies the third thoracic (T3) and first abdominal (A1) segments, whereas *abd-A* and *Abd-B* pattern the remaining abdominal segments. During early embryogenesis, the BX-C is subdivided into parasegment-specific chromosomal domains containing modular CREs that drive precise gene expression in defined posterior segments (Simon et al. 1990; Shimell et al. 2000; Qian et al. 1991; Barges et al. 2000; Zhou et al. 1999; Maeda and Karch 2006). Consistent with this organization, *abd-A* is predominantly expressed in abdominal segments A2 to A7 (Casares and Sánchez-Herrero 1995; Akam 1998), whereas *Abd-B* is expressed in all abdominal segments (A1-A7) and posterior thoracic segments during larval and adult stages (Sánchez-Herrero et al. 1985; Karch et al. 1985). In contrast, *Ubx* expression is restricted to T3 and A1 (Lewis 1978; Weatherbee et al. 1998). *BX-C* gene expression is further refined by tissue- and cell-type-specific enhancers embedded within these segment-specific regulatory domains (Simon et al. 1990; Busturia and Bienz 1993; Pirrotta et al. 1995). However, whether distinct CREs control *BX-C* gene expression in specific larval tissues, such as the fat body, remains unclear. Notably, BX-C proteins Abd-A and Abd-B are differentially expressed along the anterior-posterior axis of the larval fat body (Marchetti et al. 2003; Hemba-Waduge et al. 2025; Hemba-Waduge et al. 2026), suggesting regional heterogeneity within this tissue. Our previous work revealed that this differential heterogeneous expression defines distinct adipocyte populations with regional identities (Hemba-Waduge et al. 2025). However, the regulatory mechanisms underlying this heterogeneity, and the extent to which it is actively controlled at the transcriptional level, remain unresolved.

Adipose depots in vertebrates are similarly distributed across multiple anatomical regions and are commonly classified as subcutaneous or visceral. The species-specific placement of these depots suggests regulation by intrinsic developmental programs (Gesta et al. 2007; Parra-Peralbo et al. 2021). In mammals, adipose tissue comprises three major classes: white adipose tissue, which stores energy; brown adipose tissue, which dissipates energy through non-shivering thermogenesis; and beige (brite) adipocytes, inducible thermogenic cells that emerge within white depots in response to environmental or hormonal cues (Gesta et al. 2007; Parra-Peralbo et al. 2021; Berry et al. 2013; Sebo and Rodeheffer 2019). Molecular profiling over the past two decades has characterized gene programs underlying functional differences among these depots and revealed depot-specific *Hox* gene expression patterns (Hemba-Waduge et al. 2025).

In vertebrates, *HOX* genes are organized in four clusters, *HoxA*, *HoxB*, *HoxC*, and *HoxD*. HOX paralog groups 6-8 (*HOX6-8*) are homologous to the arthropod central Hox genes *Antp*, *Ubx*, and *abd-A*, whereas posterior *HOX9-13* genes correspond to *Abd-B* (Hubert and Wellik 2023; Mark et al. 1997; Hueber et al. 2010). These genes are differentially expressed among white and brown adipose depots, although reported patterns vary across studies, potentially reflecting differences in age, sex, or physiological state (Cantile et al. 2003; Gesta et al. 2006; Cleal et al. 2017). Notably, Clemons et al. recently demonstrated that each human adipose depot exhibits a distinct *HOX* expression signature that correlates more strongly with anatomically related tissues than with adipocyte type, supporting the idea that adipocytes may retain positional identity (Clemons 2024). Despite these associations, defining causal roles for *HOX* genes in mammalian adipose depot specification remains challenging. Functional analyses are complicated by extensive paralogy within the *HOX* gene family and by the essential roles of these genes in embryonic patterning, making it difficult to disentangle their direct effects on adipocyte metabolism from broader functions in adipogenesis and development.

Here, we identify CREs that control *abd-A* and *Abd-B* transcription specifically in abdominal adipocytes. We define a 627-bp fragment within the *Abd-B* locus and an ∼6-kb region within the *abd-A* locus that drives expression in larval adipocytes. Using the 627-bp *Abd-B* element as bait in a yeast one-hybrid (Y1H) screen, we identify three transcription factors, including Lola (Longitudinals lacking), Lolal (Lola like), and Cg (Combgap), that directly activate this CRE. Functional analyses show that Lola and Lolal regulate *Abd-B* transcription, and to a lesser extent *abd-A*, in the larval fat body. We further show that Prg (Piragua) and Svp (Seven up) are required for *Abd-B* activation. Expression of *lola*, *lolal*, *svp*, and *prg* is modulated by Wnt signaling and by Abd-B itself. Together, these results indicate that heterogeneous *Abd-B* expression in larval adipocytes is actively fine-tuned by a genetic circuit involving gene-specific CREs, multiple transcriptional factors, and Wnt signaling. In contrast, the regulatory mechanisms underlying heterogeneous *abd-A* expression are more complex and remain unresolved. These findings provide a mechanistic understanding of how the spatial patterning of *abd-A* and *Abd-B* expression influences the cellular identities across different subregions of the *Drosophila* larval fat body.

## Materials and Methods

### *Drosophila* stocks and maintenance

Flies were maintained at 25°C on a standard diet of cornmeal, molasses, and yeast. Stocks used in this study are listed in Supplementary Table S3. The *Abd.A-Gal4* line was obtained from Dr. Samir Merabet, and the *UAS-TransTimer* line from Dr. Norbert Perrimon (Hudry et al. 2011; He et al. 2019). All other stocks were obtained from the Bloomington *Drosophila* Stock Center.

### Generation of *AbdB-Gal4* and *abdA-Gal4* lines

*AbdB-Gal4* and *abdA-Gal4* driver strains were generated using a slightly modified version of the method described by Pfeiffer *et al*. (Pfeiffer et al. 2008). The enhancer region used to generate the *AbdB^GMR34G07^-Gal4* driver (1,427 bp) was subdivided into three fragments: two 400-bp fragments (*RJ1* and *RJ2*) and one 600-bp fragment (*RJ3*). The *RJ3* fragment was further subdivided into two overlapping 400-bp fragments (Fig. 4A). All fragments were PCR amplified using PrimeSTAR Max DNA polymerase (Takara, R045A) with primers containing 5’-CACC overhangs to enable directional cloning into the Gateway *pENTR/D-TOPO* vector (Thermo Fisher Scientific, K240020). LR recombination reactions (Thermo Fisher Scientific, 11791100) were used to transfer enhancer fragments into the Gateway destination vector *pBPGUw* (Addgene plasmid #17575). Following sequence validation, constructs were microinjected into *Drosophila* embryos (Rainbow Transgenic Flies and GenetiVision), and genomic integration was targeted to the *attP40* site on chromosome 2. Balanced fly stocks carrying each Gal4 driver were generated using standard genetic crosses. Genomic DNA from each line was isolated and validated by PCR and sequencing.

The *AbdB-Gal4^gypsy-RJ3-gypsy^* driver strain was generated using the *pBPGUw* vector containing the *RJ3* enhancer fragment. Two flanking gypsy insulator sequences were inserted using preamplified gypsy fragments with overlapping homology to the *pBPGUw* backbone at the *SgrDI* (Thermo Scientific, ER2031) and *NsiI* (NEB, R3127S) restriction sites. Assembly was performed using the NEBuilder HiFi DNA Assembly Cloning Kit (NEB, E5520S).

For *abdA*-Gal4 drivers, a 20 kb enhancer region was subdivided into 1.5-2.0 kb segments (Fig. 2A) and cloned following the same procedure used for *AbdB-Gal4* generation. The bacterial artificial chromosome *BACR24L18* (BACPAC Resources Center) was used as the template for generating both *abd-A* and *AbdB-Gal4* constructs. Primer sequences are listed in Table S4.

### Yeast one-hybrid (Y1H) screening

Y1H screening was performed by Hybrigenics Services (http://www.hybrigenics-services.com). The sequence for *Abd-B* (RJ3/627bp in size) fragment was PCR-amplified and cloned into the vector pB303 which was constructed by inserting the HIS3 gene into the pFL39 backbone (Bonneaud et al. 1991). Screening was performed against a random-primed *Drosophila* whole embryo cDNA library constructed into pP6 that derives from the original pGADGH plasmid (Bartel et al. 1993). 100 million clones (10-fold the complexity of the library) were screened using a mating approach with the YHGX13 (Y187 ade2-101::loxP-kanMX-loxP, matα) and Y1H300 (mata) strains as previously described (Fromont-Racine et al. 1997). 258 His+ colonies were selected on a medium lacking leucine, tryptophan and histidine and supplemented with 5 mM 3-Aminotriazol to handle bait autoactivation. The prey fragments of the positive clones were amplified by PCR and sequenced at their 5’ and 3’ junctions. The resulting sequences were used to identify the corresponding interacting proteins in the GenBank database (NCBI) using a fully automated procedure. A confidence score (PBS, for Predicted Biological Score) was attributed to each interaction as previously described (Formstecher et al. 2005; Rain et al. 2001; Wojcik et al. 2002) Twenty-one candidate proteins were identified as interacting with the 627 bp fragment. Fourteen candidates (66.7%) with moderate-confidence interaction (class D, based on global PBS scores) were excluded from downstream analysis due to potential false-positives.

### Cell biological analyses

Staining of larval fat bodies with BODIPY, DAPI, and phalloidin was performed as previously described (Zhang et al. 2017b; Li et al. 2022). Quantification for Fig. 1E, Fig. 5H, and Fig. 6H-6I were performed using ImageJ. Relative fluorescent intensity (RFI) was measured in thoracic and abdominal fat body regions using separate red and green channels. For Fig. 3M and Fig. 7G-7H, individual cells were manually delineated using the polygon selection tool in ImageJ. Cells located at tissue boundaries were excluded from analysis. Average cell size was calculated from 5-20 cells per image panel. All experiments were performed with three independent biological replicates. Statistical significance was assessed using one-tailed unpaired *t*-tests, based on a priori directional hypotheses specifying the direction of the expected outcomes.

**Fig. 1.**
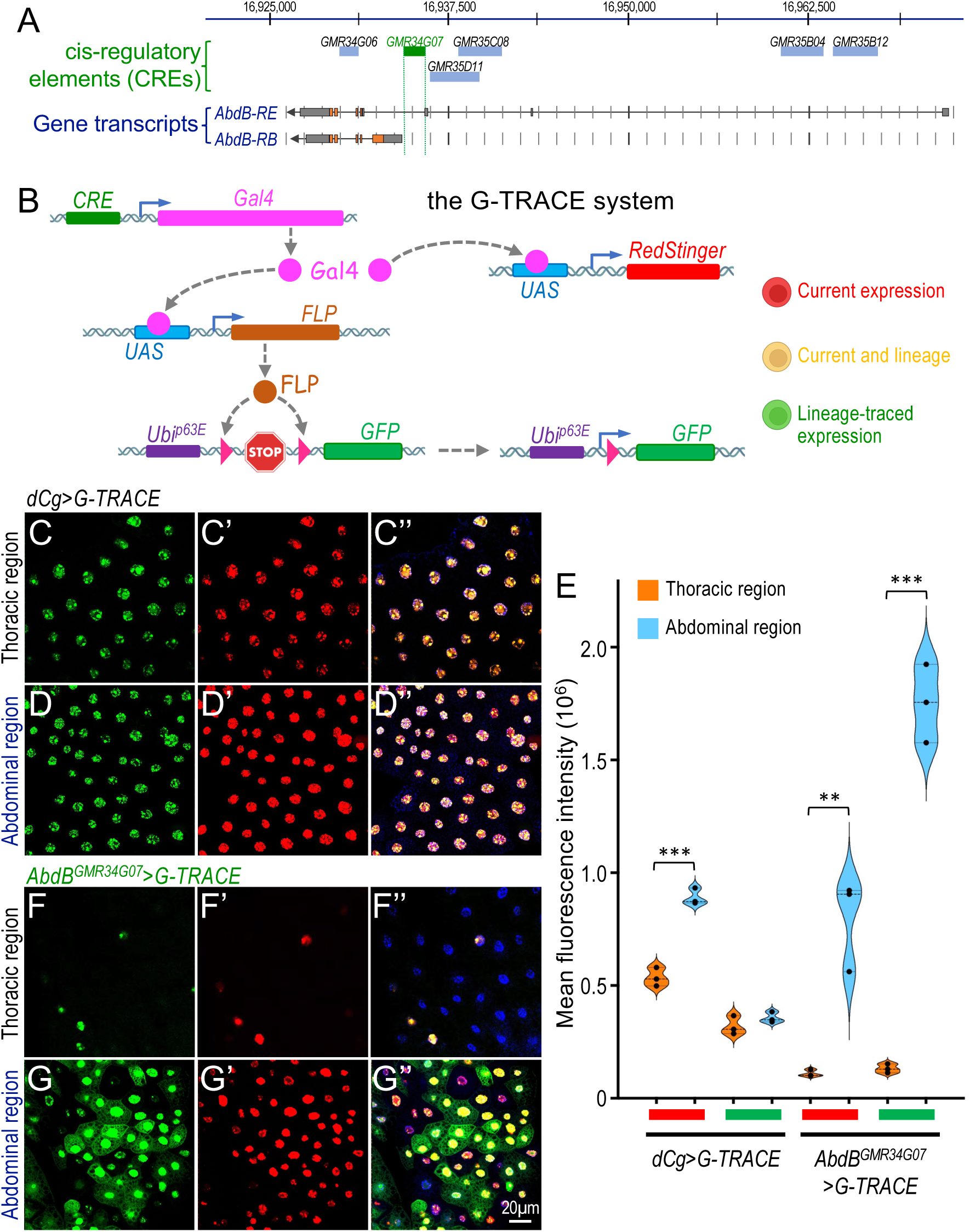
Identification of genomic regions regulating *Abd-B* expression in the larval fat body. (A) Schematic showing the relative positions of six CREs identified in *Abd-B-Gal4* driver lines. The CRE highlighted in green (GMR34G07/BL-49801) shows fat body-specific expression. Two representative *Abd-B* isoforms (RE and RB) are shown below. (B) Schematic overview of the G-TRACE system. The corresponding CRE drives Gal4 expression, activating RFP and FLP recombinase. FLP excises a STOP cassette flanked by FRT sites, removing the block between the Ubip63E promoter and nEGFP, thereby enabling EGFP expression. The real-time or current expression is marked in red, and the past or lineage-traced expression is in green. (C-G’’) Lineage-tracing using the G-TRACE system with *dCg-Gal4* and *AbdB^GMR34G07^-Gal4*. Genotypes: (C-D”) *dCg-Gal4/+; UAS-RedStinger, UAS-FLP, Ubi-p63E-STOP-GFP/+* (F-G”) *+; AbdB^GMR34G07^-Gal4/UAS-RedStinger, UAS-FLP, Ubi-p63E-STOP-GFP*. The thoracic and abdominal regions are labeled on the left side of the images. Blue indicates DAPI nuclear staining. (E) Quantification of mean fluorescence intensity of RFP and GFP signals in thoracic and abdominal regions, respectively. Signal intensity was measured across images from three independent biological replicates. Data are shown as violin plots; and genotypes are indicated below the graph. P-values were calculated using one-tailed unpaired t-tests based on directional experimental predictions. Statistical significance: P < 0.01 (**) and P < 0.001 (***). Scale bar in (G”): 20μm (applies to all images).

### The HCR RNA-FISH assay

The multiplexed *in situ* HCR RNA-FISH assay was conducted as described previously (Li et al. 2022). The following specific probe sets were obtained from Molecular Instruments: B1-Alexa Fluor 488 amplifiers with probe sets for *abd-A* (lot number PRO337); B2-Alexa Fluor 594 amplifiers with the probe sets for *Abd-B* (lot numbers PRQ416 and RTG947). Confocal images were captured using a Zeiss LSM900 confocal microscope system, and representative images for each experiment were presented.

For quantification of HCR data in larval adipocytes (Fig. 8), relative fluorescence intensity (RFI) was measured across entire cells, with background signal subtracted based on measurements from regions outside the cells. RFI quantifications for HCR experiments in adipocytes is shown in Fig. 8G-J. For HCR analyses in the VNC (Fig. S7), five independent measurements were taken per VNC to quantify the distance between *Ubx* and *Abd-B* expression domains. Three independent biological replicates were analyzed. Statistical significance was assessed using one-tailed unpaired *t*-tests, based on predefined directional hypotheses regarding the expected outcomes.

### RNA-seq analysis

RNA-seq data from previously published work (Hemba-Waduge et al. 2026) are available in the Gene Expression Omnibus (GEO) under accession number GSE280511. Newly generated RNA-seq data in this study was deposited in GEO under accession number GSE317450. RNA libraries were sequenced by BGI Americas Corporation. Sequencing reads were aligned to the *Drosophila melanogaster* dm6 reference genome using the RNASTAR aligner (Widmann et al. 2012). Gene-level count matrices were generated using featureCounts from the Subread package (Liao et al. 2014). Differential expression analysis was performed using DESeq2 (Love et al. 2014), and genes with adjusted *P* value < 0.05 were considered significantly differentially expressed. Heatmaps were generated for using the ComplexHeatmap package (Gu 2022).

### CUT&RUN assay

CUT&RUN experiments using the *dTCF^EGFP^* line together with anti-GFP antibody (Abcam, ab290) or negative control rabbit monoclonal IgG were previously described (Liu et al. 2024), and the corresponding data are available in the GEO under accession number GSE264356. CUT&RUN experiments using the *Abd.B^EGFP^* line in larval CNS were performed using the same protocol, and the resulting data was deposited in GEO under accession number GSE317446. Alignment and deduplication were performed by nf-core/cutandrun v2.0, an integrated analysis pipeline for CUT&RUN data (https://nf-co.re/cutandrun/). Downstream peak calling was performed using MACS3 with a significance threshold of q < 0.0001, followed by normalization using MACS3 (https://macs3-project.github.io/MACS/). For anti-GFP samples, alignments were provided using the “-t” option, whereas IgG controls were provided using the “-c” option. For visualization in IGV_2.14.1 (Integrative Genomics Viewer) using the *Drosophila melanogaster* dm6 genome assembly, normalized pileup bedGraph files generated by MACS3 (“-B” option) were converted to BigWig format using bedGraphicToBigWig tool from the UCSC toolkit (Liu et al. 2024).

## Results

### The identification of the genomic region responsible for the heterogeneous expression of *Abd-B* in larval fat body

To identify the cis-regulatory elements (CREs) responsible for the heterogeneous expression of *abd-A* and *Abd-B* in larval adipocytes, we screened transgenic Gal4 lines containing DNA fragments from the *BX-C* locus for their ability to drive *UAS-RedStinger* reporter expression in the larval fat body. These Gal4 lines, generated by the Rubin laboratory, include seven within the *Abd-B* locus (Table S1), but none within the *abd-A* locus (Jenett et al. 2012; Pfeiffer et al. 2008). Through this screen, we identified one Gal4 line within the *Abd-B* locus, *AbdB^GMR34G07^-Gal4* (Fig. 1A, Table S1) that exhibited specific activity in the larval fat body (see below in Fig. 1F-1G).

To further characterize the CRE activity, we used the G-TRACE (*Gal4 Technique for Real-time and Clonal Expression*) lineage tracing system, which provides spatial and temporal resolution of Gal4 activity, with nuclear EGFP labeling lineage expression and RedStinger marking current expression (Fig. 1B) (Evans et al. 2009). In this system, lineage (past) *CRE-Gal4* activity induces FLP recombinase expression, resulting in permanently GFP activation through FLP-out recombination, whereas RFP, expressed directly under UAS control, labels cells with ongoing Gal4 activity (Fig. 1B). Compared to the fat body-specific *dCg-Gal4* driver, which showed uniform expression throughout the tissue (Fig. 1C-C’’ cf. Fig. 1D-D’’; quantified in Fig. 1E), the *AbdB^GMR34G07^-Gal4* driver was specifically active in the abdominal fat body and exhibited pronounced heterogeneity among abdominal adipocytes, with substantially lower activity in the thoracic region (Fig. 1F-F’’ cf. Fig. 1G-G’’; quantified in Fig. 1E). This expression pattern was recapitulated using the dual-color fluorescent transcriptional timer reporter *UAS-TransTimer*, which combines a fast-folding destabilized GFP (edGFP) and a slow-folding, long-lived RFP to sensitively monitor dynamic transcriptional activity at single-cell resolution (Fig. S1A) (He et al. 2019). Compared to the *dCg-Gal4* driver, which displayed a uniform expression pattern throughout the larval fat body (Fig. S1B-B’’ cf. Fig. S1C-C’’), *AbdB^GMR34G07^-Gal4* exhibited significantly higher activity within the abdominal adipocytes (Fig. S1E-E’’ cf. Fig. S1D-D’’). We observed heterogeneity within the abdominal region adipocyte population, together with elevated lineage expression rather than real-time expression (Fig. S1E-E’’). Together, these observations suggest that the genomic region driving *AbdB^GMR34G07^-Gal4* expression contains a discrete CRE responsible for heterogeneous *Abd-B* expression in the abdominal fat body.

BX-C proteins have been reported to be expressed in the larval fat body by immunostaining (Marchetti et al. 2003; Duffraisse et al. 2020; Banreti et al. 2014). Using EGFP-tagged endogenous *Ubx*, *abd-A*, and *Abd-B* lines, we previously showed that Abd-A and Abd-B proteins, but not Ubx, are expressed in the larval fat body (Hemba-Waduge et al. 2026). Accordingly, our subsequent analyses focused on the regulatory mechanisms controlling the heterogeneous transcription of *abd-A* and *Abd-B* in the larval fat body.

### The identification of the genomic region responsible for the heterogeneous expression of *abd-A* in larval fat body

The expression pattern of an *abdA-Gal4* driver during embryogenesis was characterized in a previous study (Hudry et al. 2011). This *abdA-Gal4* line was inactive in the larval fat body (Fig. S2), and no other enhancer-driven Gal4 lines derived from the *abd-A* locus have been reported. Given the crucial role of Abd-A in regulating adipocyte heterogeneity (Hemba-Waduge et al. 2025) and motivated by the identification of the *AbdB^GMR34G07^-Gal4* line (Fig. 1), we employed a “promoter bashing” approach to pinpoint the regulatory region of *abd-A* responsible for its fat body-specific expression. We divided an approximately 20-kb genomic region, encompassing the *abd-A* locus, into 12 fragments, each between 1.5kb and 2.0kb. These regions were designated *JJ01* through *JJ12* (Fig. 2A). Each genomic fragment was cloned into the *pBPGUw Gal4* vector (Pfeiffer et al. 2008) and used for transgene-mediated germline transformation to generate individual fly lines reporting potential enhancer activity within *JJ01*-*JJ12*. Using the *UAS-RedStinger* reporter, we analyzed the Gal4 activities driven by each fragment and identified three out of 12 fragments (*abdA^JJ06^*, *abdA^JJ07^*, and *abdA^JJ08^*) that showed specific expression in larval fat body (Table S2).

**Fig. 2.**
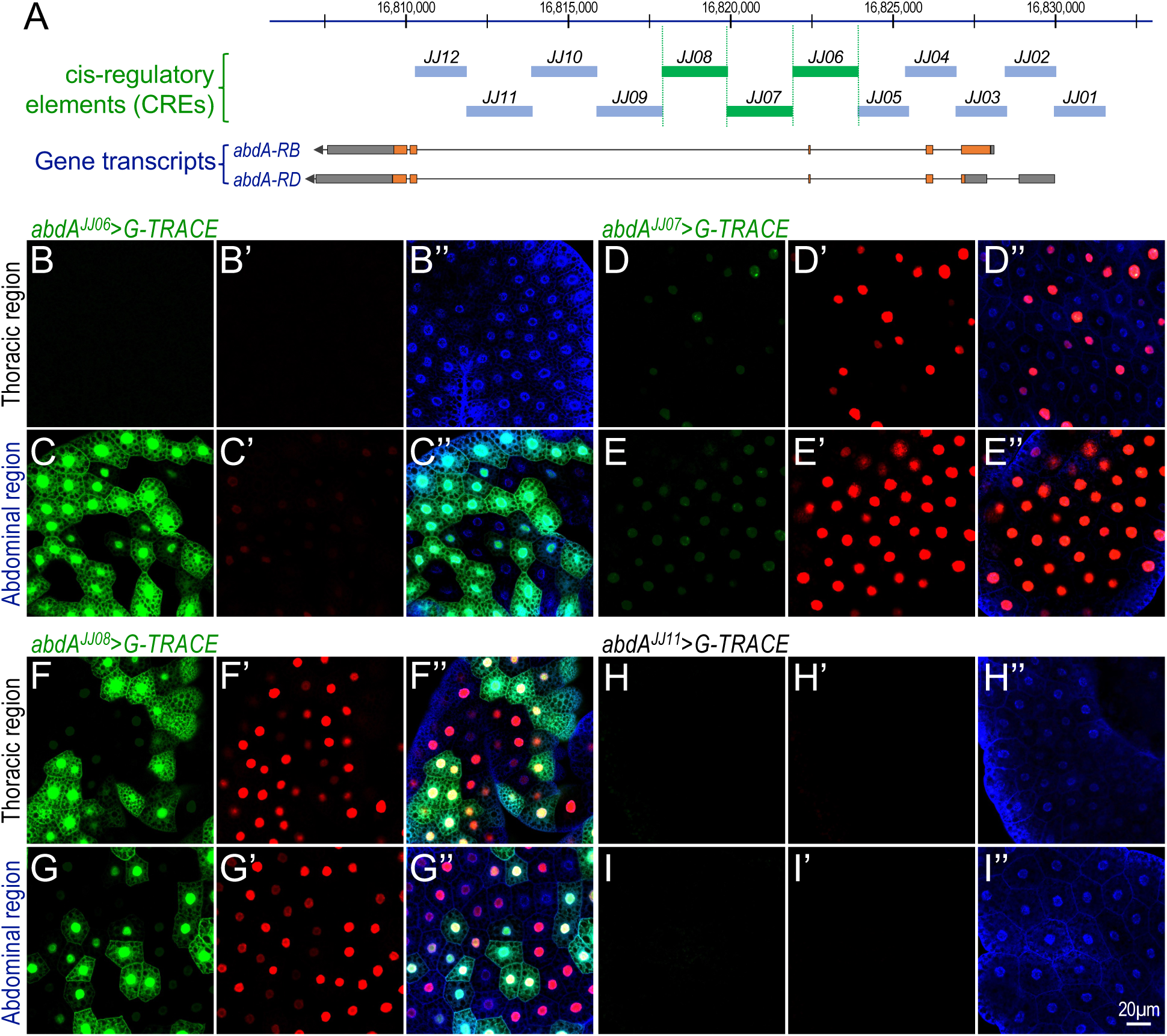
Identification of the CREs responsible for *Abd-A* expression in the larval fat body. (A) Schematic representation showing the relative positions of twelve CREs identified in *abd-A-Gal4* driver lines generated using the pBPGUw Gal4 vector. The CREs highlighted in green (*JJ06-JJ08*) exhibited distinct fat body expression. Two representative *abd-A* isoforms (RB and RD) are shown below. (B-I”) Analysis of the expression patterns of *Abd-A* (*JJ06-JJ08*) Gal4 drivers and *Abd-A^JJ11^*-Gal4 driver using G-TRACE system. Genotypes: (B-C”) *abdA^JJ06^-Gal4/+; UAS-RedStinger, UAS-FLP, Ubi-p63E-STOP-GFP/+*; (D-E”) *abdA^JJ07^-Gal4/+; UAS-RedStinger, UAS-FLP, Ubi-p63E-STOP-GFP/+*; (F-G”) *abdA^JJ08^-Gal4/+; UAS-RedStinger, UAS-FLP, Ubi-p63E-STOP-GFP/+*; and (H-I”) *abdA^JJ11^-Gal4/+; UAS-RedStinger, UAS-FLP, Ubi-p63E-STOP-GFP/+.* In panels B-I”, green indicates lineage-traced expression, red indicates transient or current expression, and blue indicates DAPI nuclear staining. The thoracic and abdominal regions are labeled on the left side of the images. Scale bar (I”): 20μm (applies to all panels).

The *abdA^JJ06^-Gal4* line exhibited region-specific expression, with strong lineage labeling (GFP-positive cells) in the abdominal fat body but no detectable current expression (RFP-positive cells) (Fig. 2C/C’). No expression was detected in the thoracic region (Fig. 2B/B’). In contrast, *abdA^JJ07^-Gal4* displayed current, but not lineage, expression in both abdominal and thoracic adipocytes (Fig. 2D/D’ and Fig. 2E/E’). The *abdA^JJ08^-Gal4* line exhibited both lineage and current activity throughout the larval fat body, with comparable expression levels in abdominal and thoracic regions (Fig. 2F/F’ and Fig. 2G/G’). Notably, although *abdA^JJ06^-Gal4* showed no detectable activity in the thoracic region (Fig. 2B/B’), both *abdA^JJ07^-Gal4* and *abdA^JJ08^-Gal4* exhibited heterogeneous expression within abdominal and thoracic adipocytes (Fig. 2D-2G). In contrast, the remaining nine lines (*abdA^JJ01^-Gal4* through *abdA^JJ05^-Gal4* and *abdA^JJ09^-Gal4* to *abdA^JJ12^-Gal4*) showed no detectable expression in the larval fat body (Fig. 2H and Fig. 2I, and Fig. S3A-E, S3I-L). Together, these observations define an approximately 6-kb CRE within the *abd-A* locus (from 3R: 16,824,029 to 16,817,958; Fig. 2A) that is sufficient to drive expression in abdominal adipocytes. These results are consistent with the endogenous expression pattern of *abd-A*, which is enriched in the abdominal larval fat body and exhibits heterogeneous expression among adipocytes (Hemba-Waduge et al. 2026).

### Depleting *Axin/Axn* using RNA interference controlled by *AbdB^GMR34G07^-Gal4* or *abdA^JJ08^-Gal4* causes severe adipocyte defects

Having identified distinct *Abd-B* and *abd-A* CREs capable of driving tissue-heterogeneous gene expression in the larval fat body, we next examined their functional contributions to adipocyte heterogeneity. As reported previously, mutation or RNAi-mediated depletion of *Axin* (*Axn*) activates Wnt/Wingless (Wg) signaling, leading to lipid mobilization and the formation of small adipocytes (Liu et al. 2024; Zhang et al. 2017a) (Fig. 3A). *Axn* encodes a core scaffold protein of the β-catenin destruction complex required for degradation of β-catenin/Armadillo (Arm) in *Drosophila* (Bejsovec 2018). We therefore used *AbdB^GMR34G07^-Gal4* and newly generated *abdA*-Gal4 drivers with larval fat body to deplete *Axn* and analyzed adipocyte heterogeneity as a functional readout.

**Fig. 3.**
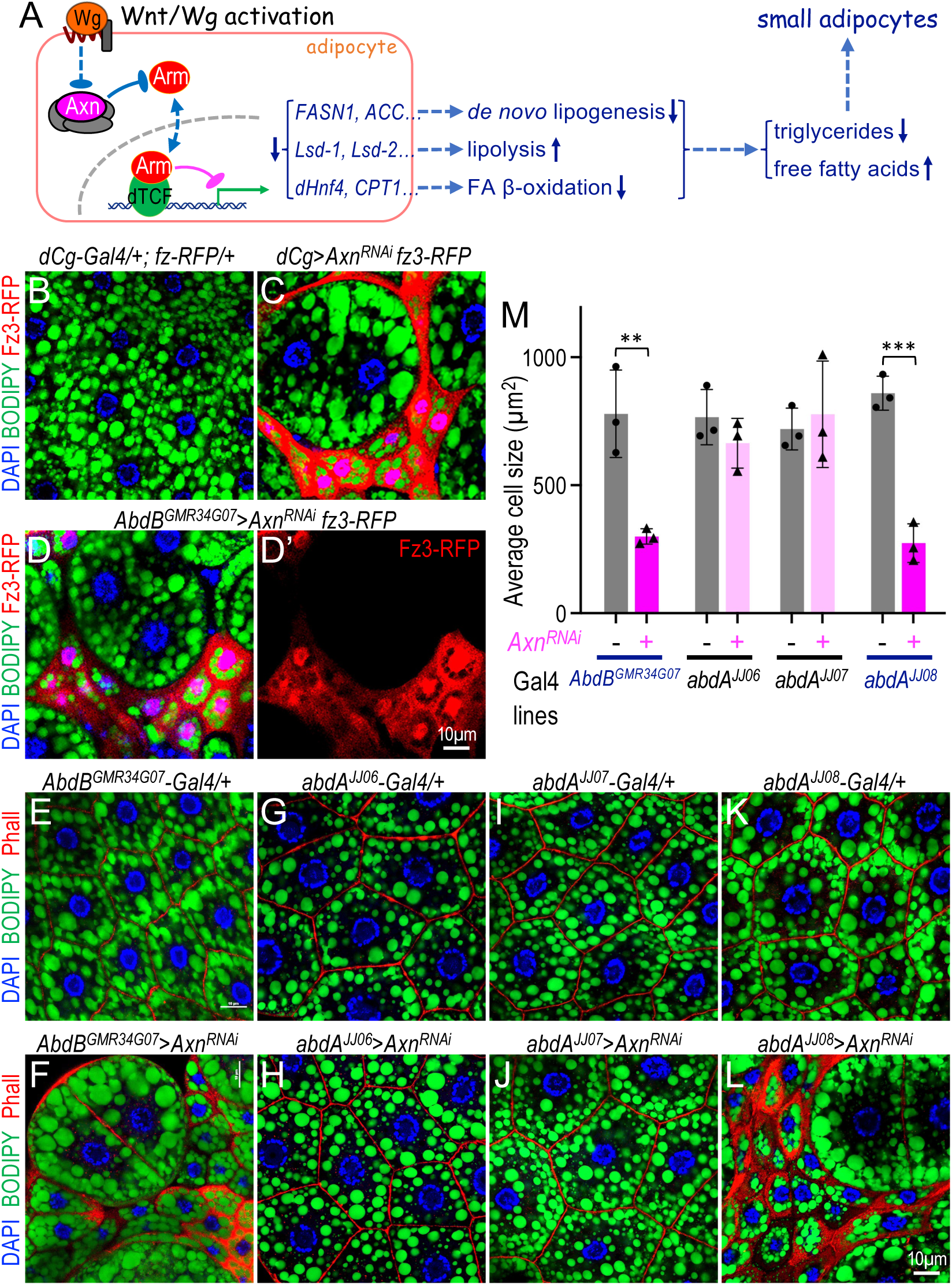
*AbdB^GMR34G07^-Gal4* and *AbdA^JJ08^-Gal4-*driven depletion of *Axn* causes severe adipocyte defects. (A) Schematic illustrating how Wnt activation in adipocytes leads to reduced triglyceride accumulation and elevated free fatty acid levels, resulting in smaller adipocytes. (B-D’) Representative confocal images of larval adipocytes stained with DAPI (blue; nuclei) and BODIPY (green; lipid droplets). *fz3*-RFP expression (red) is increased in small adipocytes in (C) and (D) but is low or absent in control adipocytes (B) and in large adipocytes. The red channel from the genotype shown in (C) is shown separately in (D’). Genotypes: (B) *dCg-Gal4/+; fz3-RFP/+*; (C) *dCg-Gal4/UAS-Axn^RNAi^; fz3-RFP/+*; and (D/D’) *UAS-Axn^RNAi^/+; AbdB^GMR34G07^/fz3-RFP.* Scale bar in (D’): 10 μm. (E-L) Representative confocal images of larval adipocytes from the indicated genotypes stained with DAPI (blue; nuclei), Phalloidin (Phall, red; cortical actin/microfilament bundles beneath the plasma membrane), and BODIPY (green; lipid droplets). Genotypes: (E) *+; AbdB^GMR34G07^-Gal4 /+*; (F) *UAS-Axn^RNAi^/+; AbdB^GMR34G07^-Gal4/+*; (G) *AbdA^JJ06^-Gal4/+; +*; (H) *UAS-Axn^RNAi^/+; AbdA^JJ06^-Gal4/+*; (I) *AbdA^JJ07^-Gal4/+; +*; (J) *UAS-Axn^RNAi^/+; AbdA^JJ07^-Gal4/+*; (K) *AbdA^JJ08^-Gal4/+; +*; and (L) *UAS-Axn^RNAi^/+; AbdA^JJ08^-Gal4/+.* Scale bar in (L): 10 μm. (M) Quantification of adipocyte size in panel E-L. Data represent three independent biological replicates, and 5-20 cells measured per replicate. Error bars indicate standard deviation. P-values were calculated using one-tailed unpaired t-tests based on directional experimental predictions. Significance levels: P < 0.01 (**) and P < 0.001 (***).

Consistent with previous findings (Hemba-Waduge et al. 2026; Liu et al. 2024), depleting *Axn* in the larval fat body using *dCg-Gal4* specifically induced the Wnt/Wg reporter *Fz3-RFP* in small adipocytes (Fig. 3C compared with control in Fig. 3B). Similarly, *Axn* depletion driven by *AbdB^GMR34G07^-Gal4* significantly increased *Fz3-RFP* expression in small adipocytes (Fig. 3D/D’), indicating that *AbdB^GMR34G07^*-positive adipocytes are associated with elevated Wnt activity. Compared to controls (Fig. 3E), *Axn* depletion using *AbdB^GMR34G07^-Gal4* produced pronounced heterogeneity among abdominal adipocytes (Fig. 3D and 3F; quantified in Fig. 3M). Likewise, *Axn* depletion driven by *abdA^JJ08^-Gal4* (Fig. 3L) induced adipocyte heterogeneity (Fig. 3L), whereas depletion using *abdA^JJ06^-Gal4* (Fig. 3H) or *abdA^JJ07^-Gal4* (Fig. 3J) did not produce detectable effects (quantified in Fig. 3M). Corresponding control genotypes (Fig. 3G/I/K) showed no detectable alterations in lipid metabolism or adipocyte morphology. The absence of phenotypes in *abdA^JJ06^-Gal4* and *abdA^JJ07^-Gal4* likely reflects their weaker lineage or real-time expression in abdominal adipocytes (Fig. 2C’ and Fig. 2E’). These observations suggest that sustained Gal4 activity, as observed with *AbdB^GMR34G07^-Gal4* and *abdA^JJ08^-Gal4*, is required to elicit detectable effects of *Axn* depletion. Together, these results indicate that the CREs asscoated with *AbdB^GMR34G07^-Gal4* and *abdA^JJ08^-Gal4* confer stronger functional activity in the larval fat body, thereby enhancing the effects of *Axn* depletion and promoting adipocyte heterogeneity relative to other *abd-A* and *Abd-B* CREs.

Although these observations reveal a correlation between *AbdB^GMR34G07^-Gal4* and *abdA^JJ08^-Gal4* activity and *Axn* depletion-induced adipocyte heterogeneity, they do not directly establish how lineage or real-time driver activity relates to Wnt-dependent regulation of lipid homeostasis at single-cell resolution. We previously showed that Wnt signaling acts together with Abd-A and Abd-B to regulate adipocyte heterogeneity in the larval fat body (Hemba-Waduge et al. 2026). Specifically, Wnt signaling promotes transcription of *abd-A* and *Abd-B*, whereas Abd-A and Abd-B function permissively to enable Wnt-mediated repression of target genes involved in lipogenesis and lipolysis (Fig. 4A) (Liu et al. 2024; Hemba-Waduge et al. 2026).

**Fig. 4.**
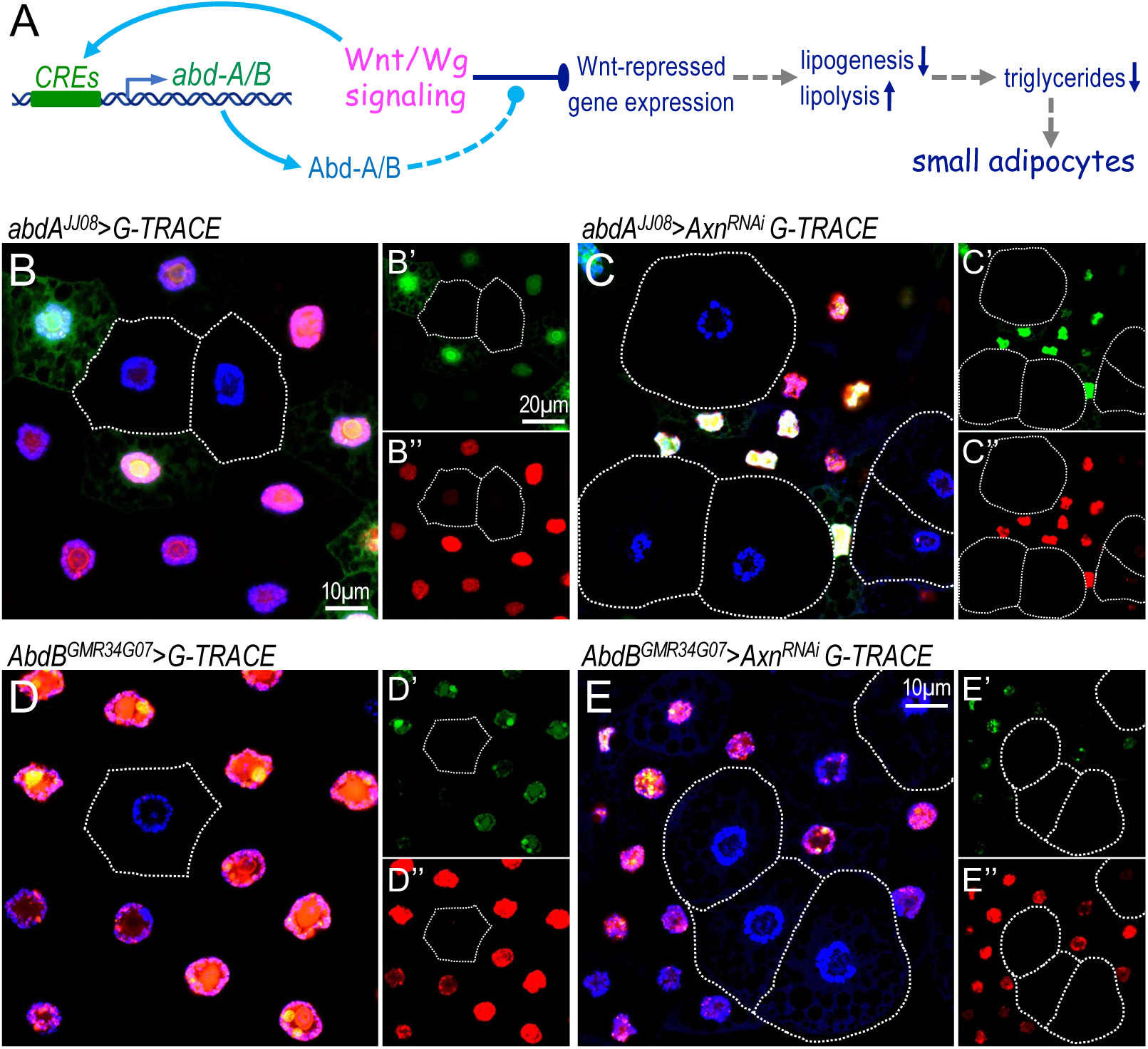
*AbdA^JJ08^-Gal4* and *AbdB^GMR34G07^-Gal4-*driven depletion of *Axn* increases *abd-A* and *Abd-B* transcription within the Wnt-induced small adipocytes. (A) Schematic illustrating how *abd-A* and *Abd-B* potentiate Wnt signaling through a feedforward loop that reinforces the interaction between Wnt signaling and *abd-A/Abd-B*. (B-E”) G-TRACE analysis following *Axn* depletion using *AbdA^JJ08^-Gal4* and *AbdB^GMR34G07^-Gal4* drivers. Genotypes: (B-B”) *AbdA^JJ08^-Gal4/+; UAS-RedStinger, UAS-FLP, Ubi-p63E-STOP-GFP/+*; (C-C”) *AbdA^JJ08^-Gal4/UAS-Axn^RNAi^; UAS-RedStinger, UAS-FLP, Ubi-p63E-STOP-GFP/+*; (D-D”) *+; AbdB-Gal4^GMR34G07^/UAS-RedStinger, UAS-FLP, Ubi-p63E-STOP-GFP*; and (E-E”) *UAS-Axn^RNAi^/+; AbdB-Gal4^GMR34G07^/UAS-RedStinger, UAS-FLP, Ubi-p63E-STOP-GFP.* Cells outlined by dotted lines in (B-B”) and (D-D”) represent adipocytes lacking *AbdA^JJ08^-Gal4* and *AbdB^GMR34G07^-Gal4* expression, respectively. Dotted outlines in (C-C”) and (E-E”) indicate large adipocytes with low Wnt activities. Green indicates lineage-traced expression, red indicates transient expression, and blue indicates DAPI nuclear staining. Scale bar in (B), 10μm (applies to panels B-E). Scale bar in (B’), 20μm (applies to panels B’/B”- E’/E”).

To determine whether *abdA^JJ08^-Gal4* and *AbdB^GMR34G07^-Gal4* activity correlates with Wnt-dependent adipocyte heterogeneity, we combined *G-TRACE* lineage analysis with Wnt activation induced by *Axn* depletion. In control animals, adipocytes positive or negative for GFP or RFP exhibited comparable cell sizes (Fig. 4B-B’’). In contrast, depletion of Axn in the *G-TRACE* background revealed a strong correlation between adipocyte size and lineage activity of *abdA^JJ08^-Gal4* (Fig. 4C-C”). Large adipocytes lacked detectable GFP or RFP signals, whereas small adipocytes exhibited strong RFP expression, either with or without GFP (Fig. 4C’/C”). These observations indicate that *abdA^JJ08^-Gal4*-active adipocytes are responsive to Wnt signaling-induced lipid mobilization, leading to reduced cell size. Similar results were obtained using *AbdB^GMR34G07^-Gal4* (Fig. 4E-E’’ compared with control in Fig. 4D-D’’).

Together, these results indicate that adipocytes with active *abd-A* and *Abd-B* expression respond to increased Wnt signaling following *Axn* depletion by reducing lipid accumulation and cell size. In contrast, adipocytes lacking *abdA^JJ08^-Gal4* and *AbdB^GMR34G07^-Gal4* activity appear refractory to Wnt-induced lipid mobilization and consequently remain enlarged, with no apparent change in lipid droplet morphology. Because Abd-A and Abd-B are required for adipocytes to respond to Wnt signaling (Hemba-Waduge et al. 2026), these observations further suggest that *abdA^JJ08^-Gal4* and *AbdB^GMR34G07^-Gal4* activity closely reflects endogenous *abd-A* and *Abd-B* expression and protein function.

### Identification of a 627bp CRE of *Abd-B* controlling adipocyte-specific expression

The robust and heterogeneous activity of the 1427 bp *Abd-B* CRE, contained within as revealed by the *AbdB^GMR34G07^-Gal4* line, prompted us to further delineate the specific elements responsible for *Abd-B* expression in the larval fat body. We divided the 1427 bp region into three fragments (Fig. 5A), each cloned into the pBPGUw vector to generate distinct Gal4 lines: *AbdB^RJ1^-Gal4*, *AbdB^RJ2^-Gal4*, and *AbdB^RJ3^-Gal4*. The activity of these potential enhancer elements was examined in larval fat body using the *UAS-RedStinger* reporter. As shown in Fig. 5C/C’ and Fig. 5D/D’, *AbdB^RJ1^-Gal4* and *AbdB^RJ2^-Gal4* exhibited no detectable Gal4 activity in the fat body, whereas *AbdB^RJ3^-Gal4* (Fig. 5E/E’) faithfully recapitulated the expression pattern of the original 1427 bp fragment (Fig. 5B/B’): *AbdB^RJ3^-Gal4* drove strong and heterogeneous expression in the abdominal region of the fat body, with little to no activity in the thoracic region, consistent with the pattern observed in the *AbdB^GMR34G07^-Gal4* line (quantified in Fig. 5H). These results indicate that the regulatory sequences mediating *AbdB^GMR34G07^-Gal4* activity in the larval fat body are localized within the 627 bp *AbdB^RJ3^* fragment.

**Fig. 5.**
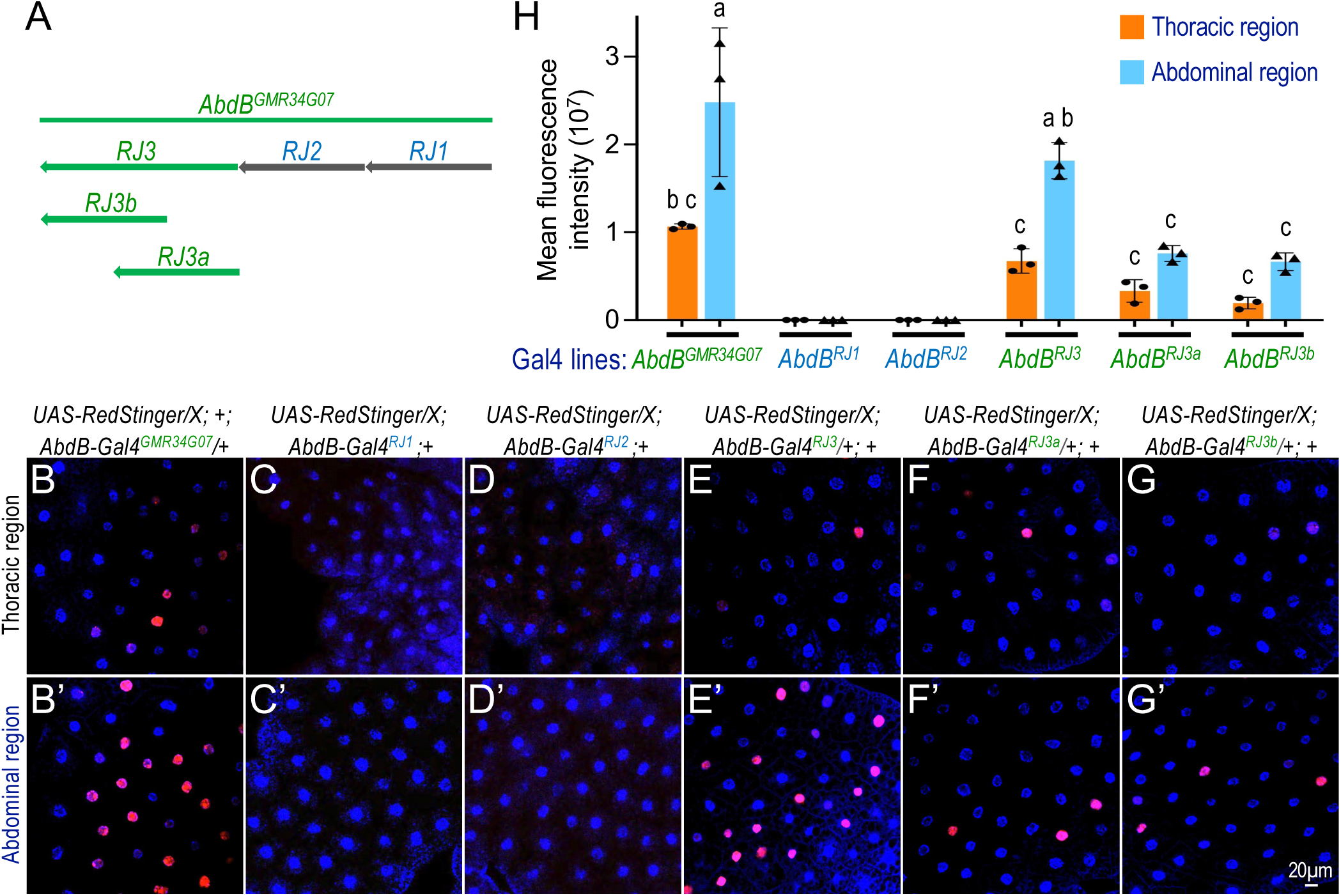
Adipocyte-specific expression of *Abd-B* is regulated by a 627bp CRE. (A) Schematic representation showing the relative positions of the *RJ1*, *RJ2* and *RJ3* fragments. The *RJ3* fragment was further subdivided into two subfragments: *RJ3a* and *RJ3b*. (B-G’) Expression patterns of newly generated *Abd-B-Gal4* drivers (*RJ1-RJ3*, *RJ3a*, and *RJ3b*) were assessed using the *UAS-RedStinger* reporter and compared with *Abd-B-Gal4^49801^.* Genotypes: (B-B’) *UAS-RedStinger/X; +; AbdB-Gal4^GMR34G07^/+*; (C-C’) *UAS-RedStinger/X; Abd.B-Gal4^RJ1^;+*; (D-D’) *UAS-RedStinger/X; Abd.B-Gal4^RJ2^;+*; (E-E’) *UAS-RedStinger/X; Abd.B-Gal4^RJ3^;+*; (F-F’) *UAS-RedStinger/X; Abd.B-Gal4^RJ3a^;+*; and (G/-G’) *UAS-RedStinger/X; Abd.B-Gal4^RJ3b^;+.* In all panels, blue indicates DAPI nuclear staining and red indicates *RedStinger* expression. The thoracic and abdominal regions are labeled on the left side of the images. The scale bar in (G’) applies to all images: 20μm. (H) Quantification of mean fluorescence intensity of the red channel within thoracic and abdominal regions. Signal intensity was measured from images obtained from three independent biological replicates. Data are presented as bar graphs, with genotypes indicated below the chart. Red represents *UAS-RedStinger* expression, and blue corresponds to DAPI nuclear staining. Statistical significance was assessed using one-way ANOVA, which identified three significantly different groups labeled a-c. Groups assigned different letters differ significantly, whereas groups sharing the same letter are not significantly different. Error bars represent standard deviation.

To further refine this CRE, we subdivided *AbdB^RJ3^* into two overlapping 400 bp fragments, *AbdB^RJ3a^*and *AbdB^RJ3b^* (Fig. 5A), and generated corresponding Gal4 driver lines (*AbdB^RJ3a^-Gal4* and *AbdB^RJ3b^-Gal4*). Using the *UAS-RedStinger* reporter, we observed that neither fragment fully reproduced the expression pattern of *AbdB^RJ3^-Gal4* or *AbdB^GMR34G07^-Gal4*: both *AbdB^RJ3a^-Gal4* and *AbdB^RJ3b^-Gal4* drove markedly weaker expression, and in fewer adipocytes (Fig. 5F/F’ and Fig. 5G/G’). These observations indicate that further subdivision of the 627-bp *AbdB^RJ3^* region disrupts CRE integrity and reduces activity, consistent with the presence of a single CRE within this sequence. Moreover, we generated another Gal4 transgenic line in which the *AbdB^RJ3^* enhancer is flanked by two *Ty3* (formerly *gypsy*) insulator sequences (*Abd.B-Gal4 ^gypsy_RJ3_^ ^gypsy^*) to minimize the effects of the surrounding chromatin environment. We observed strong and heterogeneous expression in the abdominal region of the fat body (Fig. S4A/A’), and little activity in the thoracic region (Fig. S4B/B’). Therefore, the CRE responsible for heterogeneous *Abd-B* expression in larval adipose tissue resides within the *AbdB^RJ3^* (627 bp) fragment, which drives stronger expression in abdominal adipocytes and weaker or no expression in thoracic adipocytes.

### Identification of transcription factors controlling differential heterogeneous *Abd-B* expression in the fat body

To identify the specific transcription factors that regulate *Abd-B* expression in the fat body, we performed a yeast one-hybrid (Y1H) screen (Reece-Hoyes and Marian Walhout 2012; Reece-Hoyes et al. 2011), using the 627 bp *AbdB^RJ3^* fragment as bait. We analyzed approximately 99.6 million interactions, from which 258 clones were identified as potential interactions. Among these, seven transcription factors were identified with high to moderate confidence (groups A-C in Fig. 6A), including Combgap (Cg), Charlatan (Chn), Lola (Logitudinals lacking), Lolal (Lola-like), Piragua (Prg), TBP-associated factor 1 (Taf1), and Seven up (Svp). In addition, 14 more clones exhibited moderate to low confidence interactions with the *AbdB^RJ3^*fragment.

To functionally assess the roles of these candidate factors in regulating CRE activity, we utilized the recombined *AbdB^GMR34G07^-Gal4*; *UAS-RFP* reporter (Fig. 6B/B’). If a given transcription factor affects the activity of this CRE *in vivo*, then its depletion should alter the tissue expression pattern or cell heterogeneity of RFP expression pattern driven by *AbdB^GMR34G07^-Gal4* (Fig. 6B/B’). Our analyses revealed that knockdown of *lola* (Fig. 6C/C’; multiple independent transgenic RNAi lines for *lola* in Fig. S5A-S5D) and *lolal* (Fig. 6D/D’) markedly increased RFP expression in both abdominal and thoracic adipocytes (quantified in Fig. 6H), although regional heterogeneity in the abdominal region persisted. Lola is known to regulate multiple developmental processes in *Drosophila*, including axonal guidance (Crowner et al. 2002), stem cell maintenance, and germ cell differentiation (Bass et al. 2007). *lolal* encodes a BTB/POZ domain-containing protein that interacts with Trithorax-like (Trl/GAGA associated factor), a component of Polycomb- Repressive Complexes (PRC1 and PRC2) (Faucheux et al. 2003; Mishra et al. 2003). Similarly, the depletion of *combgap* (*cg*) resulted in a pronounced increase in *Abd-B* reporter expression in abdominal adipocytes, accompanied by a modest elevation in the thoracic region (Fig. 6E/E’ cf. Fig. 6B/B’; quantified in Fig. 6H). Cg encodes a sequence-specific DNA-binding protein implicated in the recruitment of PcG proteins (Ray et al. 2016; Kassis et al. 2017), although its function in adipose tissue has not been previously characterized. These observations suggest that Lola, Lolal, and Cg transcription factors act as negative regulators of *Abd-B* promoter activity, in both thoracic and abdominal adipocytes.

In contrast, knockdown of *piragua* (*prg*) (Fig. 6F/F’) and *seven up* (*svp*) (Fig. 6G/G’) significantly reduced *Abd-B* reporter activity (quantified in Fig. 6H), indicating that both factors are required for the activation of the *AbdB^GMR34G07^* CRE. *prg* encodes a protein containing Zinc-Finger-Associated-Domain (ZAD) and C2H2 zinc-finger (ZF) domains, which are implicated in diverse developmental morphogenic processes (Nazario-Yepiz and Riesgo-Escovar 2017). Svp functions in neuronal development (Benito-Sipos et al. 2011), embryonic segmental patterning (Ponzielli et al. 2002), and insulin signaling regulation (Musselman et al. 2018; Begemann et al. 1995). To our knowledge, neither Prg nor Svp has been previously linked to gene regulation in larval adipose tissue. By contrast, depletion of *TBP-associated factor 1* (*Taf1*), *charlatan* (*chn*), and *hairy* (*hry*), three additional transcription factors identified in the Y1H screen (Fig. 6A), did not produce any detectable change in *AbdB^GMR34G07^-Gal4* activity (Fig. S5A/A’-S5C/C’), thus these transcription factors were not pursued in further analysis.

Collectively, these results identify Lola, Lolal, and Cg as negative regulators, and Prg and Svp as positive regulators, of Abd-B enhancer activity in the larval fat body. Notably, Lola and Lolal appear to play key roles in mediating the region-specific heterogeneity (thoracic vs. abdominal regions) of *AbdB^GMR34G07^-Gal4* activity.

### *Lola* and *Lolal* limit region-specific Wnt responses in adipocytes through *Abd-B* repression

As reported previously, active Wnt/Wg signaling stimulates lipid mobilization primarily in the abdominal adipocytes but not in the thoracic adipocytes (Hemba-Waduge et al. 2026). We showed that *abd-A* and *Abd-B*, which are expressed in abdominal adipocytes, act permissively to modulate Wnt target gene expression in these cells (Fig. 4A). Ectopic expression of *abd-A* or *Abd-B* in thoracic adipocytes under Wnt-activated background is sufficient to induce adipocyte defects in thoracic adipocytes (Hemba-Waduge et al. 2025; Hemba-Waduge et al. 2026). We therefore hypothesized that if depletion of *lola* or *lolal* increases *Abd-B* expression in thoracic adipocytes, then Wnt activation in this context should extend the adipocyte defects into thoracic region.

Depletion of *Axn* using the *dCg-Gal4* driver activated Wnt signaling and caused severe adipocyte defects, including a prominent heterogeneity in cell population within abdominal but not thoracic adipocytes (Fig. 7B/B’ cf. control in Fig. 7A/A’; quantified in Fig. 7G and 7H).

**Fig. 6.**
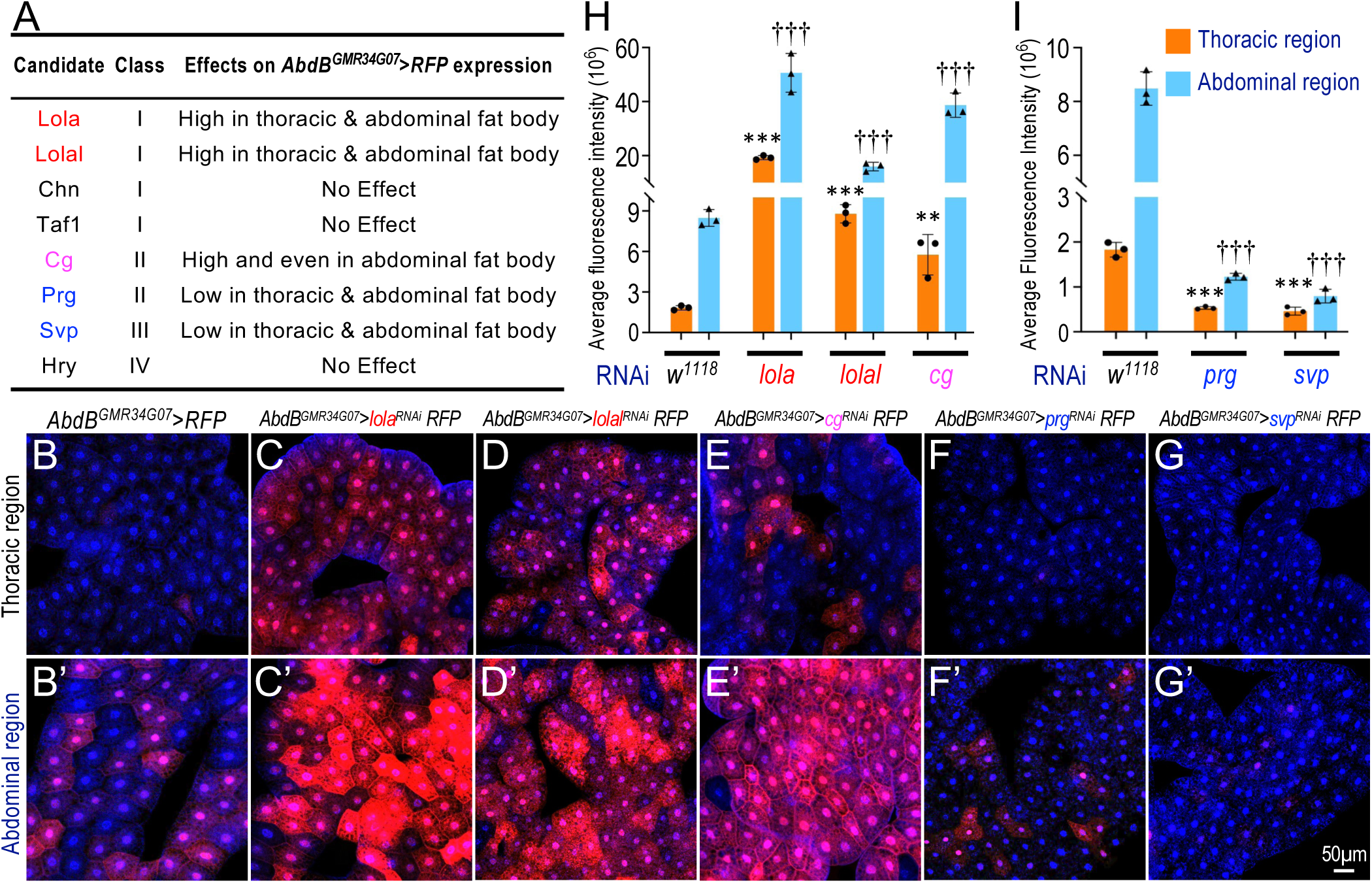
Identification of transcription factors regulating differential *Abd-B* expression in the fat body. (A) Candidate transcription factors identified from the Y1H screen were further genetically validated for their effects on *Abd-B* reporter expression. Candidate proteins were classified from ‘I’ to ‘IV’ based on the global PBS score (I: very high-confidence interaction; II: high-confidence interaction; III: good-confidence interaction; and IV: moderate-confidence interaction. (B-G’) Representative confocal images showing validation of these transcription factors using the *UAS-RFP/+; AbdB-Gal4^GMR34G07^/+* reporter. Genotypes: (B/B’) *UAS-RFP/+; AbdB-Gal4^GMR34G07^/+*; (C/C’) *UAS-RFP/+; AbdB-Gal4^GMR34G07^/UAS-lola^RNAi^*; (D/D’) *UAS-RFP/+; AbdB-Gal4^GMR34G07^/UAS-lolal^RNAi^*; (E/E’) *UAS-RFP/+; AbdB-Gal4^GMR34G07^/UAS-cg^RNAi^*; (F/F’) *UAS-RFP/+; AbdB-Gal4^GMR34G07^/UAS-prg^RNAi^*; and (G/G’) *UAS-RFP/+; AbdB-Gal4^GMR34G07^/UAS-svp^RNAi^*. In all panels, nuclei are labeled with DAPI (blue), and RFP expression is shown in red. The thoracic and abdominal regions are labeled on the left side of the images. The scale bar in G’: 50μm (applies to all images). (H, I) Quantification of the average fluorescence intensity of the red channel in the thoracic and abdominal regions. Red channel fluorescence intensity was measured from image obtained across three independent biological replicates. Data are presented as bar graphs, with the corresponding genotypes indicated below each graph. P-values were determined using one-tailed unpaired *t*-tests. Asterisks (*) indicate comparisons between the thoracic region (orange) of *dCg-Gal4/+* and the thoracic regions (blue) of the indicated genotypes, whereas daggers (†) denote comparisons between the abdominal region of *dCg-Gal4/+* with the abdominal regions of the indicated genotypes. Error bars represent standard deviations. Statistical significance is indicated as follows: P < 0.01 (**/††) and P < 0.001 (***/†††).

**Fig. 7.**
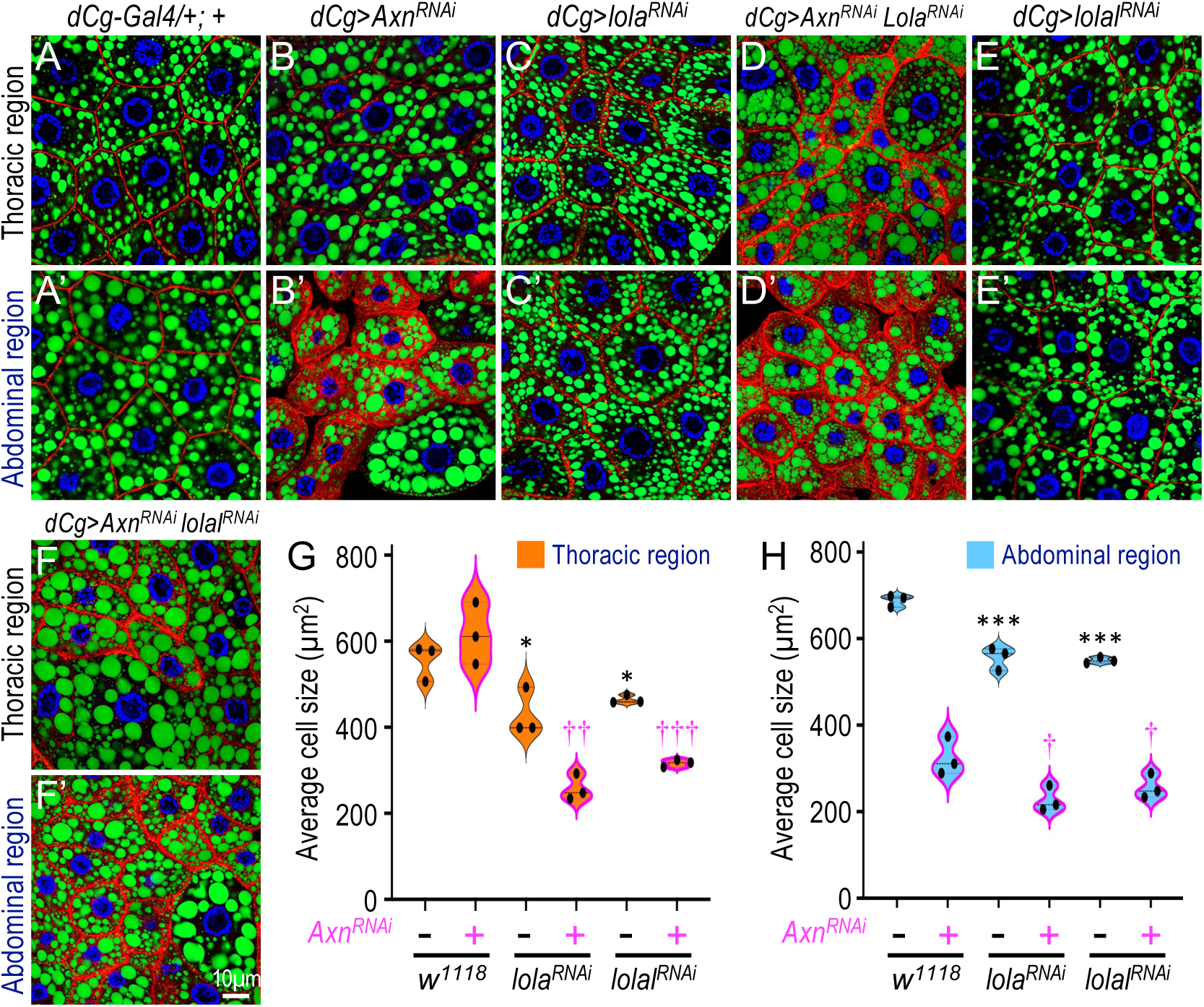
Depletion of *Lola* or *Lolal* under active Wnt signaling extends adipocyte defects into the thoracic region. (A-F’) Confocal images of larval adipocytes from the indicated genotypes stained with DAPI (blue; nuclei), Phalloidin (Phall, red; cortical microfilament bundles beneath the plasma membrane), and BODIPY (green; lipid droplets). Genotypes: (A/A’) *dCg-Gal4/+; +*; (B/B’) *dCg-Gal4/UAS-Axn^RNAi^; +*; (C/C’) *dCg-Gal4/+; UAS-lola^RNAi^/+*; (D/D’) *dCg-Gal4/ UAS-Axn^RNAi^; UAS-lola^RNAi^/+*; (E/E’) *dCg-Gal4/+; UAS-lolal^RNAi^/+*; and (F/F’) *dCg-Gal4/UAS-Axn^RNAi^; UAS-lolal^RNAi^/+.* The thoracic and abdominal regions are labeled on the left side of the images. Scale bar in panel (F’): 10μm (applies to all images). (G-H) Quantification of adipocyte size in the thoracic regions (G) corresponding to panels (A-F) and the abdominal regions (H) corresponding to panels (A’-F’). Data are presented as violin plots, with the corresponding genotypes indicated below each graph. P-values were determined using one-tailed unpaired *t*-tests based on predefined directional hypothesis for the expected outcomes. Asterisks (*) indicate comparisons between the thoracic or abdominal regions of *dCg-Gal4/+* and the corresponding regions of the other genotypes. Daggers (†) indicate comparisons between the thoracic or abdominal regions of *dCg-Gal4/UAS-Axn^RNAi^* and the corresponding regions of the other genotypes. Statistical significance is indicated as follows: P < 0.05 (*/†), P < 0.01 (**/††), and P < 0.001 (***/†††).

Consistent with our predictions, depletion of *lola* in a Wnt-activated background not only exacerbated abdominal adipocyte defects but also produced pronounced effects in thoracic adipocytes (Fig. 7D/D’; quantified in Fig. 7G and 7H). Depletion of *lola* alone did not substantially alter adipocyte morphology, aside from subtle membrane-associated changes (Fig. 7C/C’). Depletion of *lolal* under a Wnt-activated background caused qualitatively similar but weaker effects, with thoracic adipocytes showing mild heterogeneity (Fig. 7F/F’ cf. Fig. 7E/E’; quantified in Fig. 7G and 7H).

To directly assess the effects of *lola* and *lolal* depletion on *abd-A* and *Abd-B* transcription, we performed multiplexed HCR RNA-FISH (hybridization chain reaction RNA fluorescence in situ hybridization) imaging (Choi et al. 2018). Consistent with previous observations (Hemba-Waduge et al. 2026), Wnt activation caused by *Axn* depletion increased *abd-A* transcription in abdominal adipocytes, with weaker effects on *Abd-B* expression (Fig. 8B’ cf. Fig. 8A’; see Fig. S6A-D for separate channels; quantified in Fig. 8H). Thoracic adipocytes under Wnt-activated conditions showed a modest increase in *abd-A* expression compared to controls (Fig. S6A’ cf. Fig. S6B’). Depletion of *lola* significantly increased *abd-B* expression in abdominal adipocytes (Fig. 8C’ cf. Fig. 8A’) and induced a marked up-regulation of *abd-B* in thoracic adipocytes compared to controls (Fig. 8C cf. Fig. 8A; Fig. S6E and S6F; quantified in Fig. 8G). Notably, combing *lola* depletion with Wnt activation resulted in pronounced adipocyte defects in both abdominal and thoracic regions, accompanied by strong upregulation of both *abd-A* and *Abd-B* transcripts (Fig. 8D/D’ cf. Fig. 8A/A’; Fig. S6G and S6H; quantified in Fig. 8G and 8H). Under these conditions, *Abd-B* expression in abdominal adipocytes was further elevated relative to Wnt activation alone (Fig. 8D’ vs. Fig. 8A’).

**Fig. 8.**
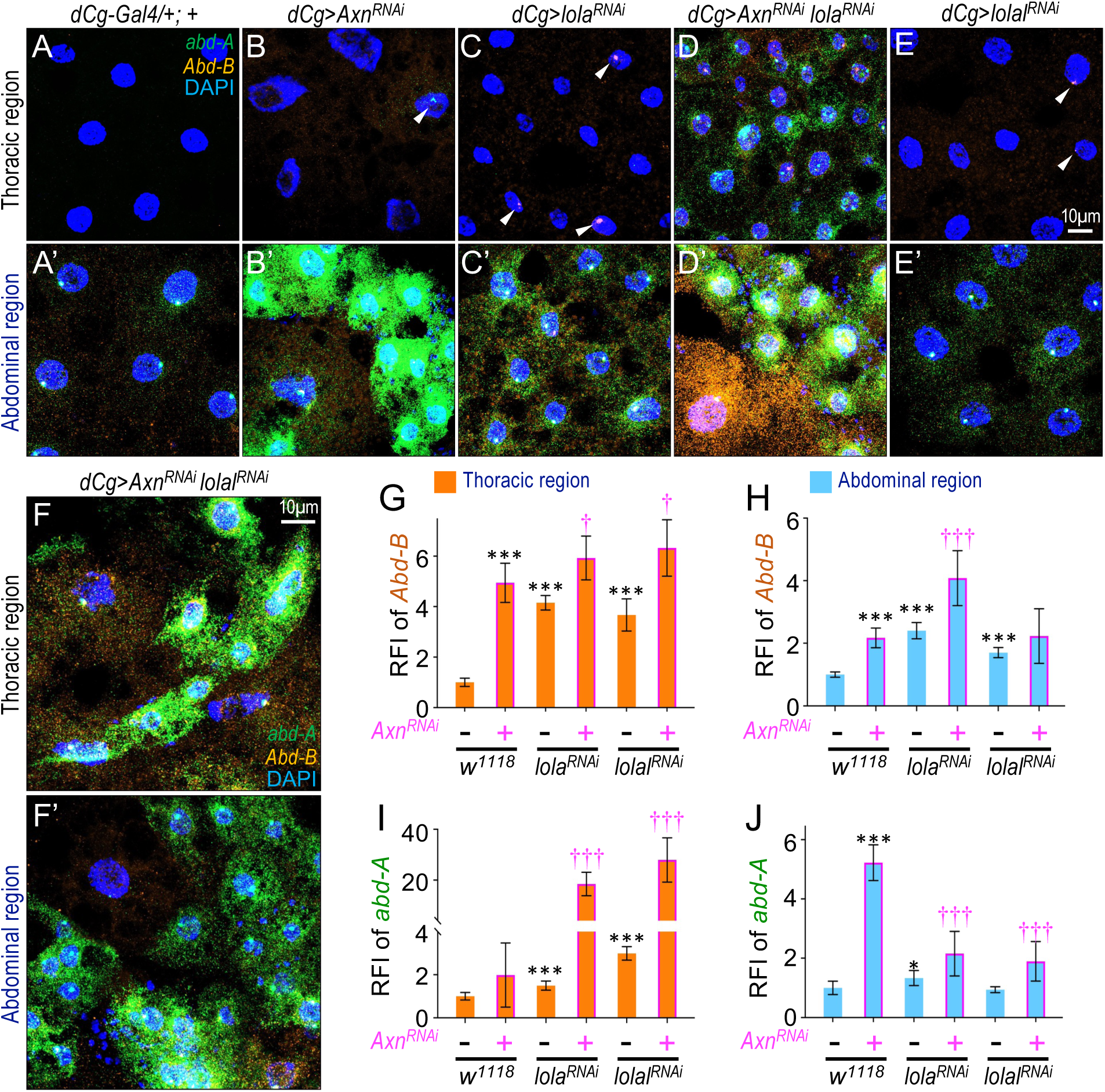
Depletion of *lola* or *lolal* under active Wnt signaling induces ectopic transcription of *abd-A* and *Abd-B* in the thoracic adipocytes. (A-F’) HCR RNA-FISH assays detecting *abd-A* (green) and *Abd-B* (orange) mRNA transcripts in larval adipocytes. White arrowheads indicate thoracic adipocytes exhibiting ectopic *abd-A* or *Abd-B* expression. Genotypes: (A/A’) *dCg-Gal4/+; +*; (B/B’) *dCg-Gal4/UAS-Axn^RNAi^; +*; (C/C’) *dCg-Gal4/+; UAS-lola^RNAi^/+*; (D/D’) *dCg-Gal4/ UAS-Axn^RNAi^; UAS-lola^RNAi^/+*; (E/E’) *dCg-Gal4/+; UAS-lolal^RNAi^/+*; and (F/F’) *dCg-Gal4/UAS-Axn^RNAi^; UAS-lolal^RNAi^/+.* The thoracic and abdominal regions are labeled on the left side of the images. The scale bar in panel (E) applies to panels from (A-E’) and represents 10μm. The scale bar in panel (F) applies to panels (F-F’) and represents 10μm. (G-J) Quantification of the relative fluorescence intensity (RFI) of *abd-A* and *Abd-B* mRNA transcripts in the thoracic regions corresponding to panels (A-F) and the abdominal regions corresponding to panels (A’-F’). The corresponding genotypes are indicated below each graph. P-values were determined using one-tailed unpaired *t*-tests. Asterisks (*) indicate comparisons between the thoracic or abdominal regions of *dCg-Gal4/+* and corresponding regions of the other genotypes. Daggers (†) indicate comparisons between the thoracic or abdominal regions of *dCg-Gal4/Axn^RNAi^*and corresponding regions of the other genotypes. Error bars represent standard deviations. Statistical significance is indicated as follows: P < 0.05 (*/†), P < 0.01 (**/††), and P < 0.001 (***/†††).

Depletion of *lolal* produced similar, but weaker effects. Transcription of *abd-B* was markedly increased in thoracic adipocytes compared to controls (Fig. 8E cf. Fig. 8A; Fig. S6I and S6J), and Wnt activation in a *lolal*-depleted background extended adipocyte defects into the thoracic region (Fig. 8F/F’ vs. Fig. 8A/A’; Fig. S6K and S6L; quantified in Fig. 8G and 8H). Taken together, these results identify Lola and Lolal as physiological repressors of *abd-B* transcription *in vivo* and reveal that they contribute to the region specificity of Wnt-dependent adipocyte responses.

### Role of Wnt signaling in potentiating *Abd-B* transcription in the larval fat body

We previously reported that active Wnt signaling potentiates, rather than activates, the expression of *abd-A* and *Abd-B* (Hemba-Waduge et al. 2026). Specifically, although *abd-A* and *Abd-B* are expressed in most abdominal adipocytes and their transcription is further enhanced by active Wnt signaling, these genes are undetectable in thoracic adipocytes. This indicates that Wnt signaling alone is insufficient to induce their *de novo* transcription (Hemba-Waduge et al. 2026), but that the severity of fat accumulation defects in the abdominal regions correlates with the degree of Wnt signaling activation. *Axn*-depletion driven by *SREBP-Gal4* produces milder phenotypes and a smaller transparent abdominal region of the larval fat body, whereas the *axn^127^* mutant causes stronger effects and a broader transparent abdominal region (Hemba-Waduge et al. 2026).

Having identified Lolal, Lola, Svp, Prg, and Cg as transcription factors responsible for directly regulating the transcription of *Abd-B*, we next asked whether their expression levels are affected by Wnt signaling. To address this, we analyzed mRNA levels of these transcription factors in control thoracic fat body samples (designated as ‘WT-Th’ for *wild-type thoracic*) and abdominal fat body samples as ‘WT-Ab’ (Hemba-Waduge et al. 2026). We observed no obvious differences between ‘WT-Th’ and WT-Ab’ samples (Fig. 9A). We then examined the effects of Wnt signaling activation by expressing an RNAi to *Axn* and analyzing thoracic (‘Axn-Th’) and abdominal (‘Axn-Ab’) fat bodies expressing *Axn^RNAi^*. As shown in Fig. 9A, Wnt activation specifically increased the mRNA levels of *lola* and *lolal*, while significantly decreasing *svp* expression in ‘Axn-Ab’ samples, but not in ‘Axn-Th’ samples. In contrast, *prg* and *cg* expression levels were not markedly affected by Wnt signaling activation (Fig. 9A).

**Fig. 9.**
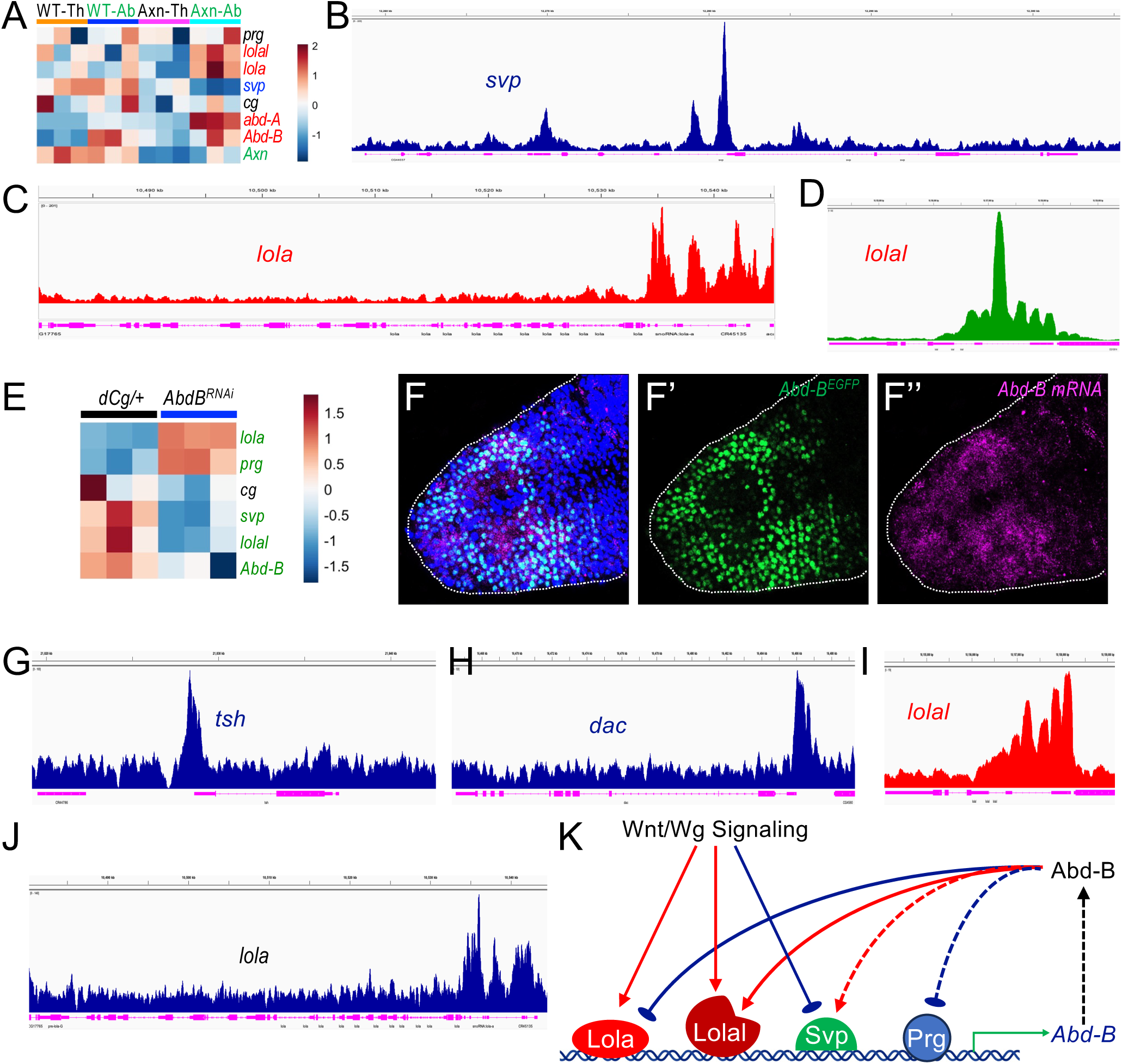
A genetic circuit regulating *Abd-B* transcription in the larval fat body. (A) Heatmap showing the effects of Wnt signaling activation on mRNA levels of five transcription factors identified from the Y1H screen. Expression levels of *abd-A*, *Abd-B*, and *Axn* are included as positive controls. WT-Th, wild-type thoracic fat body; WT-Ab, wild-type abdominal fat body; Axn-Th, thoracic fat bodies expressing *Axn^RNAi^*; Axn-Ab, abdominal fat body expressing *Axn^RNAi^*. (B-D) Genomic tracks showing dTCF binding peaks at the *svp* (B), *lola* (C), and *lolal* (D) loci in wing discs. Genes activated by Wnt signaling are shown in red, and genes inhibited by Wnt signaling are shown in blue. (E) Heatmap showing the effect of *Abd-B* depletion in fat body on mRNA levels of five transcription factors identified from the Y1H screen. *Abd-B* expression is included as a positive control. (F-F’’) Confocal images showing *Abd-B* mRNA transcripts (magenta) in the larval ventral nerve cord detected by HCR RNA-FISH together with Abd-B^EGFP^ expression (green). (F’) Single-channel image (green) showing Abd-B^EGFP^ expression, and (F”) single-channel image showing *Abd-B* mRNA transcripts (magenta) in the larval ventral nerve cord. The white dotted line marks the posterior tip of the larval ventral nerve cord. (G-J) Genomic tracks showing Abd-B binding peaks at the *tsh* (G), *dac* (H), *lolal* (I), and *lola* (J) loci in the larval CNS. (K) Model illustrating that *Abd-B* transcription in the larval fat body is regulated by multiple transcription factors, including Lola, Lolal, Svp, Prg, and Cg. Wnt signaling may further potentiate *Abd-B* expression by modulating the expression of *lola*, *lolal*, and *Svp*.

To determine whether Wnt signaling directly regulate transcription of *lola*, *lolal*, and *svp*, we analyzed previously published dTCF/Pan CUT&RUN datasets from *Drosophila* wing discs (Liu et al. 2024). dTCF/Pan is the sole transcription factor downstream of canonical Wnt/Wg signaling (Franz et al. 2017). Multiple dTCF binding peaks were detected within intronic regions of the *svp* locus (Fig. 9B), the promoter and first introns of *lola* (Fig. 9C), and the first intron of *lolal* (Fig. 9D). To validate these observations, we performed CUT&RUN targeting Pan/dTCF in larval fat body nuclei and identified multiple prominent peaks near the transcription start sites or within gene bodies of *svp*, *lola*, and *lolal* that overlapped with previously identified Pan/dTCF peaks in wing disc cells (Fig. S7). We also detected Pan/dTCF binding at both *Abd-B* and *abd-A* loci in wing discs and fat body nuclei (Fig. S8) (Hemba-Waduge et al. 2026). Although CUT&RUN from adipocyte nuclei exhibited suboptimal signal-to-noise ratio (Liu et al. 2024), the major Pan-binding sites were largely consistent across datasets.

Next, we asked whether transcription factors identified in the Y1H screen are themselves regulated by Abd-B. RNA-seq analysis of fat bodies with *Abd-B* depletion showed increased expression of *lola* and *prg*, accompanied by reduced expression of *lolal* and *svp* (Fig. 9E). Because CUT&RUN in larval fat body tissue is technically challenging (Hemba-Waduge et al. 2026; Liu et al. 2024), we performed CUT&RUN in the larval central nervous system (CNS), where Abd-B is expressed in the posterior ventral nerve cord (Fig. 9F-F’’) (Allen et al. 2020; Duckhorn 2022).

Although CNS-derived CUT&RUN may not fully recapitulate larval adipocyte chromatin states, it provides complementary and supportive evidence. To further assess regulatory relationships of *lola* and *lolal* on *Abd-B* expression in an independent context, we performed additional analyses in the larval CNS. Specifically, *lola* and *lolal* were depleted using *Elav-Gal4*, followed by HCR analysis of *abd-A* and *Abd-B* expression. Because *Elav-Gal4-*driven depletion of *lola* using resulted in lethality with the transgenic RNAi strain FBst0035721, two alternative VDRC RNAi lines were used. The larval VNC (ventral nerve cord) is well suited for these analyses, as expression domains of *abd-A*, *Abd-B*, and *Ubx* are well characterized in this tissue (Allen et al. 2020; Duckhorn 2022). As shown in Fig. S9, depleting *lola* and *lolal* in the larval CNS altered Hox domain organization in a manner consistent with effects observed in the fat body. Specifically, *Elav-Gal4-*driven knockdown reduced the distance between the *Ubx* and *Abd-B* expression domains in the VNC (Fig. S9B””-S9D”” compared to control in Fig. S9A””; quantified in Fig. S9E), indicating an anterior expansion of the *Abd-B* domain. These findings parallel our fat body data, where depleting *lola* and *lolal* in a Wnt-activated background expanded Wnt-induced adipocyte phenotypes into more anterior (thoracic) regions. Together, these results support a role for *lola* and *lolal* in restricting *Abd-B* expression boundaries across multiple larval tissues.

Abd-B has previously been reported to negatively regulate *teashirt* (*tsh*) and *dachshund* (*dac*) in posterior tissues across different developmental contexts, although whether these interactions are direct remains unclear (Wang et al. 2013; Estrada and Sanchez-Herrero 2001). Using CUT&RUN profiling using the *Abd-B^EGFP^* line, we identified at least one prominent binding peak within each of the *tsh* and *dac* loci (Fig. 9G,H), serving as positive controls and supporting the possibility that Abd-B directly regulates these targets. CUT&RUN analysis of larval CNS further revealed significant Abd-B occupancy at the *lolal* (Fig. 9I), and *lola* loci (Fig. 9J), suggesting potential direct regulation these transcription factors. In contrast, genome browser tracks at the *svp*, *prg*, and *cg* loci did not meet peak-calling thresholds in our analyses (Fig. S10).

Collectively, RNA-seq and CUT&RUN data support a regulatory model in which *Abd-B* expression is positively regulated by Lola, Lolal, and Cg, and negatively regulated by Svp and Prg (Fig. 6). Conversely, *lola* and *prg* are negatively regulated by Abd-B, whereas *lolal* and *svp* are positively regulated by Abd-B, together forming a feedback regulatory network. This circuit is further modulated by Wnt signaling, which promotes *lola* and *lolal* transcription while repressing *svp* expression (Fig. 9A). Finally, CUT&RUN analyses of dTCF/Pan and Abd-B indicate that regulation of *lola*, *lolal*, and *svp* is likely direct, as evidenced by the presence of binding peaks for both factors at these loci (Fig. 9K).

## Discussion

Hox proteins encoded in the *Bithorax Complex*, including Abd-A and Abd-B, are central determinants of posterior segment identity in *Drosophila*. We previously showed that differential expression of endogenous *abd-A* and *Abd-B* in larval fat bodies defines adipocyte heterogeneity (Hemba-Waduge et al. 2026; Hemba-Waduge et al. 2025). However, how this heterogeneity is transcriptionally established has remained unresolved. Here, we identify cis-regulatory elements (CREs) and transcription factors that, together with Wnt signaling, form a genetic circuit controlling heterogeneous *Abd-B* expression in larval adipocytes (Fig. 9K). These findings provide mechanistic insight into how spatially patterned *Hox* gene expression contributes to the cellular identity of subregions of the *Drosophila* larval fat body.

### *cis*-regulatory control of *Abd-B* transcription in larval adipocytes

By mapping CREs within the *Abd-B* locus, we identified a 627 bp region within *Abd-B* that drives region-specific and heterogeneous expression in the larval fat body. This enhancer activity is consistent with our previous transcriptomic analyses showing enriched *Abd-B* expression in abdominal relative to thoracic adipocytes. Y1H screen and functional genetic analyses further revealed that multiple transcription factors, including Lola, Lolal, and Svp, modulate both reporter activity and endogenous *Abd-B*, indicating that *Abd-B* transcription is controlled by a combinatorial regulatory network in the larval fat body (Fig. 9K).

Lola and Lolal are paralogous BTB (BR-C, ttk and bab) or POZ (Pox virus and Zinc finger) domain-containing proteins that share high sequence similarity but have diverged functionally (Zollman et al. 1994; Bardwell and Treisman 1994; Costoya 2007; Hu et al. 2021; Hu et al. 2011). Lola has been implicated in axon guidance and ovarian cell death (Crowner et al. 2002; Bass et al. 2007), whereas Lolal functions as an effector of the Trl-GAF (Trithorax-like/GAFA factor)-mediated Polycomb Group (PcG) regulation (Faucheux et al. 2003; Mishra et al. 2003). Our data suggest a distinct role for these paralogs in larval adipocytes: Lolal is required for proper regulation of endogenous *Abd-B* expression, whereas Lola contributes primarily to maintaining transcriptional difference between thoracic and abdominal fat body regions.

Lolal (also known as Batman) has been shown to act in both activation and repression of homeotic genes, including repression of *Sex combs reduced* (*Scr*) (Faucheux et al. 2003). Through its BTB/POZ domain, Lolal directly interacts with Trl/GAF protein, a factor implicated in PcG recruitment and chromatin organization (Mishra et al. 2003). Lolal also genetically interacts with Polyhomeotic (Ph), and both are required for PRE (Polycomb Response Element)-mediated silencing (Faucheux et al. 2003; Mishra et al. 2003), suggesting that Lolal may facilitate PcG-dependent repression at the *Abd-B* locus. In addition, recent work has shown that Trl/GAF can promote long-range chromatin interactions via its POZ/BTB domain (Li et al. 2023), raising the possibility that Lolal contributes to high-order chromatin looping between *Abd-B* promoters and distal enhancers.

In addition to Lola and Lolal, Y1H screening identified Cg, Svp and Prg as direct binders of the 627 bp *Abd-B* enhancer fragment. Functional analysis using the *AbdB^GMR34G07^-Gal4* reporter revealed distinct regulatory effects: depletion of *cg* increased reporter activity, whereas depletion of *svp* and *prg* reduced it. Like Lolal, Cg is also involved in recruiting PcG proteins (Kassis et al. 2017). Notably, depletion of Cg resulted in more uniform *AbdB^GMR34G07^-Gal4* reporter expression within the abdominal fat body together with comparatively milder effects within the thoracic region, suggesting a role for Cg in generating interregional heterogeneity. In contrast, depleting either Lola or Lolal expanded reporter expression into thoracic regions while preserving heterogeneity, indicating that these factors primarily regulate the regional identity of larval adipocytes. Further evidence suggested that depleting *lola* and *lolal* under an activated Wnt signaling regime extended the adipocyte defects into the thoracic region (Fig. 7D and Fig. 7F cf. Fig. 7B), potentially driven by increased *Abd-B* transcription within thoracic adipocytes (Fig. 8D and Fig. 8F cf. Fig. 8B). This aligns with our previous observations, in which we have demonstrated that ectopic expression of Abd-B in a Wnt-activated regime is sufficient to cause adipocyte defects in the thoracic region (Hemba-Waduge et al. 2026).

Together, our findings support a model in which heterogeneous *Abd-B* transcription in larval adipocytes emerges from the integration of enhancer-specific transcription factor inputs and higher-order chromatin organization. Long-range promoter-enhancer interactions, potentially mediated by BTB/POZ domain-containing proteins, may enable precise spatial control of *Abd-B* transcription. The incomplete overlap between reporter activity and endogenous expression likely reflects the modular and hierarchical regulatory architecture of the *Bithorax Complex*. Indeed, Abd-B expression is controlled by multiple parasegment-specific regulatory domains separated by boundary elements that both restrict enhancer crosstalk and permit long-distance regulatory interactions (Postika et al. 2018). Tissue-specific outputs further depend on clustered transcription factor binding sites within cis-regulatory modules (Starr et al. 2011). Thus, while *AbdB^GMR34G07^-Gal4* serves as a useful qualitative proxy for *Abd-B* transcriptional activity, it captures only a subset of the regulatory logic governing endogenous *Abd-B* expression.

### Establishing transcription factor-binding site relationships

Many eukaryotic transcription factors recognize short DNA motifs that occur frequently throughout the genome (Ekker et al. 1991; Choi and Sinha 2006). However, only a subset of these sites is functionally engaged in transcriptional regulation. Specific gene regulation is achieved, in part, through the combinatorial context of neighboring transcription factor-binding sites and through protein-protein interactions among bound factors. Consequently, identifying and characterizing clusters of transcription factor-binding sites is essential for defining the rules that govern binding specificity and for determining the regulatory outcomes of transcription factor-DNA complex formation, including transcriptional activation or repression. Characterization of CREs, such as those identified here, is therefore critical for decoding the mechanisms underlying complex transcriptional control of key target genes.

### Transcriptional regulation of *abd-A* in larval fat bodies

In contrast to the better-defined mechanisms regulating heterogeneous *Abd-B* transcription, the transcriptional control of *abd-A* in larval adipocytes remains poorly understood. Using a promoter-bashing approach, we identified an ∼6-kb CRE within the *abd-A* locus (*abdA^JJ06^*, *abdA^JJ07^*, and *abdA^JJ08^*) that drives region-specific and heterogeneous expression in the larval fat body. Within this region, the ∼2-kb fragment (*abdA^JJ08^*) most closely recapitulates endogenous *abd-A* expression, as its activity is sufficient to drive adipocyte heterogeneity upon *Axn* depletion (Fig. 3). Notably, G-TRACE analyses suggest that *abdA^JJ06^* is active in early adipocyte lineages but show low activity in third-instar larval fat bodies, whereas *abdA^JJ07^* exhibited opposite pattern. In contrast, *abdA^JJ08^* is active in both lineage tracing and third-instar larval fat body (Fig. 2), potentially providing a sufficient temporal window for effective RNAi-mediated *Axn* depletion by this CRE (Fig. 3).

Despite these advances, we were unable to further delineate the minimal regulatory elements controlling *abd-A* expression in larval adipocytes, highlighting the complexity of its regulatory architecture. Notably, depletion of *lola* and *lolal* produced comparable effects on *abd-A* and *Abd-B* expression (Fig. 8), suggesting that these *Hox* genes share partially overlapping regulatory inputs while also being subject to gene-specific control mechanisms. Further comparative analyses of the three CREs (*abdA^JJ06^*, *abdA^JJ07^*, and *abdA^JJ08^*) may therefore provide an entry point to dissect the regulatory logic underlying heterogeneous *abd-A* transcription in larval fat bodies. Given the essential role of *abd-A*, and its major contribution to adipocyte heterogeneity (Hemba-Waduge et al. 2026; Hemba-Waduge et al. 2025), elucidating the regulatory mechanisms governing its transcription remains an important objective for future studies.

### A genetic circuit regulating *Abd-B* transcription in larval fat bodies

Our previous work showed that *abd-A* and *Abd-B* expression is potentiated by active Wnt/Wg signaling, while Abd-A and Abd-B themselves act permissively to enable Wnt target gene expression (Fig. 4A) (Hemba-Waduge et al. 2026). Although the mechanistic basis of this reciprocal relationship remains unresolved, our current findings suggest that Wnt signaling promotes *Abd-B* transcription through a transcriptional regulatory circuit summarized in Fig. 9K. This model integrates genetic, transcriptomic, and chromatin-binding data to explain how Wnt/Wg signaling further potentiates spatially heterogeneous *Abd-B* expression. While key components of this circuit are supported by Abd-B CUT&RUN data generated from larval central nervous tissue, Abd-B binding may be tissue-specific, and future chromatin profiling in larval adipocytes will be required to fully validate this model.

Consistent with this framework, CUT&RUN analysis in the larval central nervous tissue revealed Abd-B occupancy at multiple nodes of the proposed regulatory network, including the *svp*, *lola*, and *lolal* loci, indicating that Abd-B may directly participate in shaping its own transcriptional landscape. Integration of these binding data with RNA-seq and functional perturbation analyses supports a model in which *Abd-B* expression is governed by a balance of activating and repressive inputs. Lola, Lolal, and Cg function as positive regulators of *Abd-B* transcription, whereas Svp and Prg provide inhibitory inputs. In turn, Abd-B feeds back onto this network by repressing the expression of *lola* and *prg* while activating *lolal* and *svp*, thereby establishing interconnected feedback loops that constrain *Abd-B* expression in a spatially heterogeneous manner. Wnt/Wg signaling further biases this circuit toward *Abd-B* activation by promoting *lola* and *lolal* expression while repressing *svp*. Supporting direct regulation, CUT&RUN profiles for both dTCF/Pan and Abd-B show binding within the *lola*, *lolal*, and *svp* loci (Fig. 9K), indicating that Wnt signaling and Abd-B converge on shared transcriptional targets to stabilize patterned *Abd-B* expression.

Taken together, this study advances our understanding of the regulatory mechanisms controlling *BX-C* gene expression in larval adipocytes. By delineating the relevant CREs and transcription factors, it highlights the complexity of postembryonic *BX-C* gene regulation. The observed heterogeneity in *abd-A* and *Abd-B* expression, as well as regional differences in larval fat body, suggest a potential role for these *Hox* genes in regulating adipocyte development and metabolic homeostasis. Future studies will be essential to define how Lolal and Lola modulate heterogeneous *abd-A* and *Abd-B* transcription and to determine the functional consequences for adipocyte heterogeneity and lipid metabolism.

In higher organisms, adipose tissues are distributed across multiple anatomical depots defined by intrinsic developmental programs. Extensive studies have characterized depot-specific gene expression under diverse physiological and pathological conditions, revealing molecular bases for functional heterogeneity. However, depot-specific expression of *Hox* gene expression and its functional consequences remain poorly understood. Although several studies have reported depot-restricted *Hox* expression in mice and humans, results vary across depots and studies (see recent review (Hemba-Waduge et al. 2025)), highlighting a complex, context-dependent regulatory landscape. Given the high evolutionary conservation of the transcription factors identified here from *Drosophila* to humans, these regulatory mechanisms may be broadly conserved.

### Limitations of the study

In addition to the limitations discussed above, our study has several constraints. First, although our data implicate CREs and transcription factor networks in controlling heterogeneous *abd-A* and *Abd-B* expression, we lack direct evidence for enhancer-promoter interactions or higher-order chromatin looping. Analysis of chromatin architecture in larval adipocytes remains technically challenging due to polyploid, limited cell numbers, and abundant lipid droplets, which currently preclude high-resolution chromosome conformation assays such as Micro-C. Second, Polycomb group (PcG) and Trithorax group (TrxG) complexes are classically considered key regulators for maintaining postembryonic Hox gene expression (Kassis et al. 2017; Schuettengruber et al. 2017; Beck et al. 2010; Kuroda et al. 2020). However, our findings suggest that *BX-C* gene regulation may be active and compensatory rather than passive maintenance, with tissue-specific mechanisms involving novel regulators such as Lola and Lolal predominating in this context. Third, our proposed regulatory model (Fig. 9K) is derived primarily from genetic perturbations and bulk RNA-seq analyses, which capture steady-state transcriptional outputs rather than dynamic regulatory behavior. Given that the circuit includes multiple feedforward and feedback interactions, time-resolved and quantitative approaches, including mathematical modeling, will be required to fully understand how heterogeneous *abd-A* and *Abd-B* expression is established and maintained in the larval fat body.

## Data availability

*Drosphila* strains generated in this study are available upon request. Supplemental material available at GENETICS online.

## Acknowledgments

We thank Samir Merabet and Norbert Perrimon for generously providing fly strains, Tzu-Hao Liu and Jasmine Sun for technical assistance, and Tianyi Zhang for insightful comments on the manuscript. We also thank the Bloomington *Drosophila* Stock Center (NIH Grant P40OD018537) for providing fly stocks.

## Funding

This work was supported by the National Institute of Health (GM129266 to J.-Y.J.).

## Conflicts of interest

The authors declare no competing interests.

## Author contributions

R.-U.-S. H.-W., and J.-Y.J. designed the experiments; R.-U.-S. H.-W., M.L., X.L., and E.A.B. conducted the experiments with input from S.E.B and K.A.M.; R.-U.-S. H.-W., M.L., X.L., S.E.B, K.A.M., and J.-Y.J. analyzed the data; R.-U.-S. H.-W., and J.-Y.J. wrote the manuscript. All authors reviewed and revised the manuscript.

## Figure legends

**Fig. S1 Validation of CREs responsible for *Abd-B* expression in the larval fat body.** (A) Schematic diagram illustrating the major steps underlying the UAS-TransTimer system. Upon CRE-driven Gal4 expression, the “TransTimer” cassette, containing destabilized GFP (dGFP) and stable RFP, is activated. In this system, current/real-time expression is marked by green fluorescence, whereas lineage or past expression is indicated by red fluorescence. To further analyze the activity of the *dCg-Gal4* and *AbdB^GMR34G07^-Gal4* drivers, we employed the *UAS-TransTimer* reporter system. Genotypes: (B-C’’) *dCg-Gal4/+; pUAST-edGFP::2A::RFP/+.* (D-E’’) *+; pUAST-edGFP::2A::RFP/AbdB^GMR34G07^-Gal4.* (B-B” and D-D”) thoracic region; (C-C” and E-E”) abdominal region. Green indicates transient or current expression, red indicates lineage or past expression, and blue corresponds to DAPI staining marking nuclei. Scale bar in panel C: 20μm (applies to all images).

**Fig. S2 Expression pattern of *abdA-Gal4* in the larval fat body.** Lineage tracing analysis of the *abdA-Gal4* driver using the G-TRACE system. Genotype: *+; abdA-Gal4/UAS-RedStinger, UAS-FLP,Ubi-p63E(^FRT^STOP^FRT^)GFP.* Red fluorescence represents *RedStinger* expression, indicating current Gal4 activity, whereas green fluorescence indicates GFP expression, marking lineage-tracing cells. (A-A”) thoracic region; (B-B”) abdominal region. Scale bar in panel A: 20μm (applies to all images).

**Fig. S3 Expression patterns of *AbdA-Gal4* drivers generated using the pBPGUw-Gal4 vector.** The *UAS-RedStinger* reporter system was used to examine the expression patterns of the *abdA-Gal4* drivers. Genotypes: (A/A’) *UAS-RedStinger/X; abdA^JJ01^-Gal4/+*; (B/B’) *UAS-RedStinger/X; abdA^JJ02^-Gal4/+*; (C/C’) *UAS-RedStinger/X; abdA^JJ03^-Gal4/+*; (D/D’) *UAS-RedStinger/X; abdA^JJ04^-Gal4//+*; (E/E’) *UAS-RedStinger/X; abdA^JJ05^-Gal4/+*; (F/F’) *UAS-RedStinger/X; abdA^JJ06^-Gal4/+*; (G/G’) *UAS-RedStinger/X; abdA^JJ07^-Gal4/+*; (H/H’) *UAS-RedStinger/X; abdA^JJ08^-Gal4/+*; (I/I’) *UAS-RedStinger/X; abdA^JJ09^-Gal4/+*; (J/J’) *UAS-RedStinger/X; abdA^JJ10^-Gal4/+*; (K/K’) *UAS-RedStinger/X; abdA^JJ11^-Gal4/+*; and (L/L’) *UAS-RedStinger/X; abdA^JJ12^-Gal4/+.* The thoracic and abdominal regions are labeled on the left side of the images. SG: Salivary glands. Scale bar in panel A: 50μm (applies to all images).

**Fig. S4 The *Abd-B* cis-regulatory element *RJ3* flanked by gypsy insulators exhibits region-specific expression patterns similar to those of *RJ3* without gypsy insulators.** The *UAS-RedStinger* reporter system was used to examine the expression pattern of the *abdB-Gal4* driver containing the RJ3 fragment flanked by gypsy insulators. Genotype: *UAS-RedStinger/X; AbdB-Gal4^gypsy-RJ3-gypsy^/+; +.* Red fluorescence indicates RedStinger expression driven by the corresponding *AbdB*-Gal4 construct. Scale bar in panel B: 20μm (applies to all images).

**Fig. S5 Validation of additional Y1H assay candidates using the *UAS-RFP* combined with the *AbdB-Gal4^GMR34G07^* driver.** (A-D’) The effects of depleting *lola* were further validated using two additional RNAi lines from the Vienna *Drosophila* Resource Center (VDRC: #101925 and #12573). The *UAS-RFP/+; AbdB-Gal4^GMR34G07^/+* reporter system was used. In all panels, blue indicates DAPI staining marking nuclei, and red indicates RFP expression. Scale bar in panel D’: 50μm (applies to all images in A-D’). Genotypes: (A/A’) *UAS-RFP/+; AbdB-Gal4^GMR34G07^/+*; (B/B’) *UAS-RFP/+; AbdB-Gal4^GMR34G07^/UAS-lola^RNAi^*; (C/C’) *UAS-RFP/+; AbdB-Gal4^GMR34G07^/UAS-lola^RNAi^ [VDRC# 101925]*; *and* (D/D’) *UAS-RFP/+; AbdB-Gal4^GMR34G07^/UAS-lola^RNAi^ [VDRC# 12573].* (E-G’) Analysis of *Taf1*, *chn*, and *hry* depletion using the *UAS-RFP* recombined with the *AbdB-Gal4^GMR34G07^* driver. Genotypes: (E/E’) *UAS-RFP/+; AbdB-Gal4^GMR34G07^/UAS-Taf1^RNAi^*; (F/F’) *UAS-RFP/+; AbdB-Gal4^GMR34G07^/UAS-Chn^RNAi^*; and (G/G’) *UAS-RFP/+; AbdB-Gal4^GMR34G07^/UAS-Hry^RNAi^*. The thoracic and abdominal regions are labeled on the left side of the images. Scale bar in panel G’: 50 μm (applies to images E-G’).

**Fig. S6: Analysis of *abd-A* and *Abd-B* transcript levels following depletion of *lola* or *lolal* under conditions of activated Wnt signaling.** HCR RNA-FISH was used to detect *abd-A* (green) and *Abd-B* (orange) mRNA transcripts in larval adipocytes. White arrowheads indicate thoracic adipocytes expressing *abd-A* or *Abd-B* transcripts. Genotypes: (A-A’’) and (B-B’’) *dCg-Gal4/+; +*; (C-C’’) and (D-D’’) *dCg-Gal4/UAS-Axn^RNAi^; +*; (E-E’’) and (F-F’’) *dCg-Gal4/+; UAS-lola^RNAi^/+*; (G-G’’) and (H-H’’) *dCg-Gal4/UAS-Axn^RNAi^; UAS-lola^RNAi^/+*; (I-I’) and (J-J’’) *dCg-Gal4/+; UAS-lolal^RNAi^/+*; and (K-K’’) and (L-L’’) *dCg-Gal4/ UAS-Axn^RNAi^; UAS-lolal^RNAi^/+.* The thoracic and abdominal regions are labeled on the left side of the images. Scale bar in panel A: 10μm (applies to all images).

**Fig. S7: Genome browser tracks displaying dTCF/Pan binding peaks at the *svp* (A), *lola* (B), and *lolal* (C) loci.** Peaks were identified by CUT&RUN analysis performed on wing imaginal discs (upper panels, dark blue indicating downregulated expression and red indicating upregulated expression upon Wnt/Wg activation) and purified adipocyte nuclei from larval fat bodies (lower panels, orange). Tracks are visualized in the IGV_2.14.1 browser. The y-axis is autoscaled, and four gene isoforms per locus are displayed in magenta. Shared prominent peaks between datasets are indicated by arrows. (*) indicates transcription start sites.

**Fig. S8: Genome browser tracks showing dTCF/Pan binding peaks at the *Abd-B* (A) and *abd-A* (B) loci.** Peaks were identified by CUT&RUN analysis performed on wing imaginal discs (upper panels, red indicating upregulated expression upon Wnt/Wg activation) and purified adipocyte nuclei from larval fat bodies (lower panels, orange). Tracks are visualized in the IGV_2.14.1 browser. The y-axis is autoscaled, and four gene isoforms per locus are displayed in magenta. Shared prominent peaks between datasets are indicated by arrows. (*) indicates transcription start sites. Scale bars: 2.5 kb.

**Fig. S9: *Elav-Gal4*-driven depletion of *lola* or *lolal* markedly expands the *Abd-B* expression domain in the larval ventral nerve cord (VNC).** HCR RNA-FISH was used to detect *abd-A* (green), *Abd-B* (orange), and *Ubx* (magenta) mRNA transcripts in the larval VNC. Scale bar in panel D: 10μm (applies to all images). Genotypes: (A-A””) *UAS-RFP/+; AbdB-Gal4^GMR34G07^/+*; (B-B””) *UAS-RFP/+; AbdB-Gal4^GMR34G07^/UAS-lolal^RNAi^*; (C-C””) *UAS-RFP/+; AbdB-Gal4^GMR34G07^/UAS-lola^RNAi^ [VDRC #101925]*; and (D-D””) *UAS-RFP/+; AbdB-Gal4^GMR34G07^/UAS-lola^RNAi^ [VDRC #12573].* (E) Quantification of the normalized gap between *Abd-B* and *Ubx* expression domains in the VNC (n = 3 independent biological replicates, with 5 independent measurements obtained from each VNC). P values were calculated using one-tailed unpaired *t*-tests, based on predefined directional predictions in the experimental design. Error bars represent standard deviations. ***: P<0.001.

**Fig. S10: Genome browser tracks showing Abd-B binding peaks at the *svp* (A), *prg* (B), and *cg* (C) loci.** Peaks were identified by CUT&RUN analysis performed on larval central nervous system (CNS), with enrichment primarily in the ventral nerve cord (VNC). Tracks are visualized in the IGV_2.14.1 browser. The y-axis is autoscaled. (*) indicates transcription start sites.

## Notes

### Competing Interest Statement

The authors have declared no competing interest.

## Literature cited

Akam, M., 1998 Hox genes, homeosis and the evolution of segment identity: no need for hopeless monsters. Int J Dev Biol 42 (3):445–451.

Allen, A.M., M.C. Neville, S. Birtles, V. Croset, C.D. Treiber et al., 2020 A single-cell transcriptomic atlas of the adult Drosophila ventral nerve cord. Elife 9.

Banreti, A., B. Hudry, M. Sass, A.J. Saurin, and Y. Graba, 2014 Hox proteins mediate developmental and environmental control of autophagy. Dev Cell 28 (1):56–69.

Bardwell, V.J., and R. Treisman, 1994 The POZ domain: a conserved protein-protein interaction motif. Genes & Development 8 (14):1664–1677.

Barges, S., J. Mihaly, M. Galloni, K. Hagstrom, M. Müller et al., 2000 The *Fab-8* boundary defines the distal limit of the bithorax complex *iab-7* domain and insulates *iab-7* from initiation elements and a PRE in the adjacent *iab-8* domain. Development 127 (4):779-790.

Bartel, P., C.T. Chien, R. Sternglanz, and S. Fields, 1993 Elimination of false positives that arise in using the two-hybrid system. Biotechniques 14 (6):920–924.

Bass, B.P., K. Cullen, and K. McCall, 2007 The axon guidance gene lola is required for programmed cell death in the Drosophila ovary. Dev Biol 304 (2):771–785.

Beck, S., F. Faradji, H. Brock, and F. Peronnet, 2010 Maintenance of Hox gene expression patterns. Adv Exp Med Biol 689:41–62.

Begemann, G., A.-M. Michon, L. V.D. Voorn, R. Wepf, and M. Mlodzik, 1995 The *Drosophila* orphan nuclear receptor Seven-up requires the Ras pathway for its function in photoreceptor determination. Development 121 (1):225–235.

Bejsovec, A., 2018 Wingless Signaling: A Genetic Journey from Morphogenesis to Metastasis. Genetics 208 (4):1311–1336.

Benito-Sipos, J., C. Ulvklo, H. Gabilondo, M. Baumgardt, A. Angel et al., 2011 Seven up acts as a temporal factor during two different stages of neuroblast 5-6 development. Development 138 (24):5311–5320.

Berry, D.C., D. Stenesen, D. Zeve, and J.M. Graff, 2013 The developmental origins of adipose tissue. Development 140 (19):3939–3949.

Bonneaud, N., O. Ozier-Kalogeropoulos, G.Y. Li, M. Labouesse, L. Minvielle-Sebastia et al., 1991 A family of low and high copy replicative, integrative and single-stranded S. cerevisiae/E. coli shuttle vectors. Yeast 7 (6):609–615.

Busturia, A., and M. Bienz, 1993 Silencers in abdominal-B, a homeotic Drosophila gene. The EMBO Journal 12 (4):1415–1425.

Cantile, M., A. Procino, M. D’Armiento, L. Cindolo, and C. Cillo, 2003 HOX gene network is involved in the transcriptional regulation of in vivo human adipogenesis. J Cell Physiol 194 (2):225–236.

Casares, F., and E. Sánchez-Herrero, 1995 Regulation of the *infraabdominal* regions of the bithorax complex of *Drosophila* by gap genes. Development 121 (6):1855–1866.

Choi, H.M.T., M. Schwarzkopf, M.E. Fornace, A. Acharya, G. Artavanis et al., 2018 Third-generation in situ hybridization chain reaction: multiplexed, quantitative, sensitive, versatile, robust. Development 145 (12).

Choi, Y.S., and S. Sinha, 2006 Determination of the consensus DNA-binding sequence and a transcriptional activation domain for ESE-2. Biochem J 398 (3):497–507.

Cleal, L., T. Aldea, and Y.Y. Chau, 2017 Fifty shades of white: Understanding heterogeneity in white adipose stem cells. Adipocyte 6 (3):205–216.

Clemons, H.J., Hogan, D.J., Brown, P.O., 2024 Depot-Specific mRNA Expression Programs in Human Adipocytes Suggest Physiological Specialization via Distinct Developmental Programs. bioRxiv.

Costoya, J.A., 2007 Functional analysis of the role of POK transcriptional repressors. Briefings in Functional Genomics and Proteomics 6 (1):8–18.

Crowner, D., K. Madden, S. Goeke, and E. Giniger, 2002 Lola regulates midline crossing of CNS axons in *Drosophila*. Development 129 (6):1317–1325.

Diaz-Cuadros, M., O. Pourquié, and E. El-Sherif, 2021 Patterning with clocks and genetic cascades: Segmentation and regionalization of vertebrate versus insect body plans. PLOS Genetics 17 (10):e1009812.

Duckhorn, J.C., Junker, I.P., Ding, Y., Shirangi, T.R., 2022 Combined in Situ Hybridization Chain Reaction and Immunostaining to Visualize Gene Expression in Whole-Mount Drosophila Central Nervous Systems, in Behavioral Neurogenetics. Neuromethods, edited by D. Yamamoto. Humana, New York, NY.

Duffraisse, M., R. Paul, J. Carnesecchi, B. Hudry, A. Banreti et al., 2020 Role of a versatile peptide motif controlling Hox nuclear export and autophagy in the Drosophila fat body. J Cell Sci 133 (18).

Ekker, S.C., K.E. Young, D.P. von Kessler, and P.A. Beachy, 1991 Optimal DNA sequence recognition by the Ultrabithorax homeodomain of Drosophila. EMBO J 10 (5):1179–1186.

Estrada, B., and E. Sanchez-Herrero, 2001 The Hox gene Abdominal-B antagonizes appendage development in the genital disc of Drosophila. Development 128 (3):331–339.

Evans, C.J., J.M. Olson, K.T. Ngo, E. Kim, N.E. Lee et al., 2009 G-TRACE: rapid Gal4-based cell lineage analysis in Drosophila. Nature Methods 6 (8):603–605.

Faucheux, M., J.Y. Roignant, S. Netter, J. Charollais, C. Antoniewski et al., 2003 batman Interacts with polycomb and trithorax group genes and encodes a BTB/POZ protein that is included in a complex containing GAGA factor. Mol Cell Biol 23 (4):1181–1195.

Formstecher, E., S. Aresta, V. Collura, A. Hamburger, A. Meil et al., 2005 Protein interaction mapping: a Drosophila case study. Genome Res 15 (3):376–384.

Franz, A., D. Shlyueva, E. Brunner, A. Stark, and K. Basler, 2017 Probing the canonicity of the Wnt/Wingless signaling pathway. PLoS Genet 13 (4):e1006700.

Fromont-Racine, M., J.C. Rain, and P. Legrain, 1997 Toward a functional analysis of the yeast genome through exhaustive two-hybrid screens. Nat Genet 16 (3):277–282.

Gesta, S., M. Bluher, Y. Yamamoto, A.W. Norris, J. Berndt et al., 2006 Evidence for a role of developmental genes in the origin of obesity and body fat distribution. Proc Natl Acad Sci U S A 103 (17):6676–6681.

Gesta, S., Y.H. Tseng, and C.R. Kahn, 2007 Developmental origin of fat: tracking obesity to its source. Cell 131 (2):242–256.

Gu, Z., 2022 Complex heatmap visualization. iMeta 1 (3):e43.

He, L., R. Binari, J. Huang, J. Falo-Sanjuan, and N. Perrimon, 2019 In vivo study of gene expression with an enhanced dual-color fluorescent transcriptional timer. Elife 8.

Hemba-Waduge, R.U., M. Liu, and J.Y. Ji, 2025 Transcriptional Regulation of Lipid Metabolism by Wnt Signaling and Hox Protein Cues. Adv Exp Med Biol 1482:61–81.

Hemba-Waduge, R.U., M. Liu, X. Li, J.L. Sun, E.A. Budslick et al., 2026 Adipocyte heterogeneity regulated by the Bithorax Complex-Wnt signaling crosstalk in Drosophila. EMBO Reports 27:367–386.

Hu, Y., A. Comjean, J. Rodiger, Y. Liu, Y. Gao et al., 2021 FlyRNAi.org—the database of the Drosophila RNAi screening center and transgenic RNAi project: 2021 update. Nucleic Acids Research 49 (D1):D908–D915.

Hu, Y., I. Flockhart, A. Vinayagam, C. Bergwitz, B. Berger et al., 2011 An integrative approach to ortholog prediction for disease-focused and other functional studies. BMC Bioinformatics 12 (1):357.

Hubert, K.A., and D.M. Wellik, 2023 Hox genes in development and beyond. Development 150 (1).

Hudry, B., S. Viala, Y. Graba, and S. Merabet, 2011 Visualization of protein interactions in living Drosophila embryos by the bimolecular fluorescence complementation assay. BMC Biology 9 (1):5.

Hueber, S.D., G.F. Weiller, M.A. Djordjevic, and T. Frickey, 2010 Improving Hox protein classification across the major model organisms. PLoS One 5 (5):e10820.

Jaeger, J., 2011 The gap gene network. Cellular and Molecular Life Sciences 68 (2):243–274.

Jenett, A., G.M. Rubin, T.T. Ngo, D. Shepherd, C. Murphy et al., 2012 A GAL4-driver line resource for Drosophila neurobiology. Cell Rep 2 (4):991–1001.

Karch, F., B. Weiffenbach, M. Peifer, W. Bender, I. Duncan et al., 1985 The abdominal region of the bithorax complex. Cell 43 (1):81–96.

Kassis, J.A., J.A. Kennison, and J.W. Tamkun, 2017 Polycomb and Trithorax Group Genes in Drosophila. Genetics 206 (4):1699–1725.

Kimelman, D., and B.L. Martin, 2012 Anterior–posterior patterning in early development: three strategies. WIREs Developmental Biology 1 (2):253–266.

Kuroda, M.I., H. Kang, S. De, and J.A. Kassis, 2020 Dynamic Competition of Polycomb and Trithorax in Transcriptional Programming. Annu Rev Biochem 89:235–253.

Lewis, E.B., 1978 A gene complex controlling segmentation in Drosophila. Nature 276 (5688):565–570.

Li, X., X. Tang, X. Bing, C. Catalano, T. Li et al., 2023 GAGA-associated factor fosters loop formation in the Drosophila genome. Mol Cell 83 (9):1519–1526.e1514.

Li, X., M. Zhang, M. Liu, T.H. Liu, R.U. Hemba-Waduge et al., 2022 Cdk8 attenuates lipogenesis by inhibiting SREBP-dependent transcription in Drosophila. Dis Model Mech 15 (11):dmm049650.

Liao, Y., G.K. Smyth, and W. Shi, 2014 featureCounts: an efficient general purpose program for assigning sequence reads to genomic features. Bioinformatics 30 (7):923–930.

Liu, M., R.U. Hemba-Waduge, X. Li, X. Huang, T.H. Liu et al., 2024 Wnt/Wingless signaling promotes lipid mobilization through signal-induced transcriptional repression. Proc Natl Acad Sci U S A 121 (28):e2322066121.

Love, M.I., W. Huber, and S. Anders, 2014 Moderated estimation of fold change and dispersion for RNA-seq data with DESeq2. Genome Biol 15 (12):550.

Maeda, R.K., and F. Karch, 2006 The ABC of the BX-C: the bithorax complex explained. Development 133 (8):1413–1422.

Marchetti, M., L. Fanti, M. Berloco, and S. Pimpinelli, 2003 Differential expression of the Drosophila BX-C in polytene chromosomes in cells of larval fat bodies: a cytological approach to identifying in vivo targets of the homeotic Ubx, Abd-A and Abd-B proteins. Development 130 (16):3683–3689.

Mark, M., F.M. Rijli, and P. Chambon, 1997 Homeobox genes in embryogenesis and pathogenesis. Pediatr Res 42 (4):421–429.

Martin, A.C., 2020 The Physical Mechanisms of *Drosophila* Gastrulation: Mesoderm and Endoderm Invagination. Genetics 214 (3):543–560.

Martin, A.C., M. Kaschube, and E.F. Wieschaus, 2009 Pulsed contractions of an actin–myosin network drive apical constriction. Nature 457 (7228):495–499.

Mishra, K., V.S. Chopra, A. Srinivasan, and R.K. Mishra, 2003 Trl-GAGA directly interacts with lola like and both are part of the repressive complex of Polycomb group of genes. Mech Dev 120 (6):681–689.

Musselman, L.P., J.L. Fink, E.J. Maier, J.A. Gatto, M.R. Brent et al., 2018 Seven-Up Is a Novel Regulator of Insulin Signaling. Genetics 208 (4):1643–1656.

Nazario-Yepiz, N.O., and J.R. Riesgo-Escovar, 2017 piragua encodes a zinc finger protein required for development in Drosophila. Mech Dev 144 (Pt B):171–181.

Parra-Peralbo, E., A. Talamillo, and R. Barrio, 2021 Origin and Development of the Adipose Tissue, a Key Organ in Physiology and Disease. Front Cell Dev Biol 9:786129.

Pfeiffer, B.D., A. Jenett, A.S. Hammonds, T.T. Ngo, S. Misra et al., 2008 Tools for neuroanatomy and neurogenetics in Drosophila. Proc Natl Acad Sci U S A 105 (28):9715–9720.

Pirrotta, V., C.S. Chan, D. McCabe, and S. Qian, 1995 Distinct parasegmental and imaginal enhancers and the establishment of the expression pattern of the Ubx gene. Genetics 141 (4):1439–1450.

Ponzielli, R., M. Astier, A. Chartier, A. Gallet, P. ThéRond et al., 2002 Heart tube patterning in *Drosophila*requires integration of axial and segmental information provided by the *Bithorax Complex*genes and *hedgehog*signaling. Development 129 (19):4509–4521.

Postika, N., M. Metzler, M. Affolter, M. Müller, P. Schedl et al., 2018 Boundaries mediate long-distance interactions between enhancers and promoters in the Drosophila Bithorax complex. PLoS Genet 14 (12):e1007702.

Qian, S., M. Capovilla, and V. Pirrotta, 1991 The bx region enhancer, a distant cis-control element of the Drosophila Ubx gene and its regulation by hunchback and other segmentation genes. The EMBO Journal 10 (6):1415–1425.

Rain, J.C., L. Selig, H. De Reuse, V. Battaglia, C. Reverdy, et al., 2001 The protein-protein interaction map of Helicobacter pylori. Nature 409 (6817):211–215.

Ray, P., S. De, A. Mitra, K. Bezstarosti, J.A.A. Demmers et al., 2016 Combgap contributes to recruitment of Polycomb group proteins in *Drosophila*. Proceedings of the National Academy of Sciences 113 (14):3826–3831.

Reece-Hoyes, J.S., A. Diallo, B. Lajoie, A. Kent, S. Shrestha et al., 2011 Enhanced yeast one-hybrid assays for high-throughput gene-centered regulatory network mapping. Nat Methods 8 (12):1059–1064.

Reece-Hoyes, J.S., and A.J. Marian Walhout, 2012 Yeast one-hybrid assays: a historical and technical perspective. Methods 57 (4):441–447.

Saunders, T.E., 2021 The early Drosophila embryo as a model system for quantitative biology. Cells Dev 168:203722.

Schuettengruber, B., H.M. Bourbon, L. Di Croce, and G. Cavalli, 2017 Genome Regulation by Polycomb and Trithorax: 70 Years and Counting. Cell 171 (1):34–57.

Sebo, Z.L., and M.S. Rodeheffer, 2019 Assembling the adipose organ: adipocyte lineage segregation and adipogenesis in vivo. Development 146 (7).

Shimell, M.J., A.J. Peterson, J. Burr, J.A. Simon, and M.B. O’Connor, 2000 Functional analysis of repressor binding sites in the iab-2 regulatory region of the abdominal-A homeotic gene. Dev Biol 218 (1):38–52.

Simon, J., M. Peifer, W. Bender, and M. O’Connor, 1990 Regulatory elements of the bithorax complex that control expression along the anterior-posterior axis. The EMBO Journal 9 (12):3945–3956.

Soshnikova, N., and D. Duboule, 2009 Epigenetic Temporal Control of Mouse Hox Genes in Vivo. Science 324 (5932):1320-1323.

Starr, M.O., M.C. Ho, E.J. Gunther, Y.K. Tu, A.S. Shur et al., 2011 Molecular dissection of cis-regulatory modules at the Drosophila bithorax complex reveals critical transcription factor signature motifs. Dev Biol 359 (2):290–302.

Struhl, G., 1981 A homoeotic mutation transforming leg to antenna in Drosophila. Nature 292 (5824):635–638.

Sánchez-Herrero, E., I. Vernós, R. Marco, and G. Morata, 1985 Genetic organization of Drosophila bithorax complex. Nature 313 (5998):108–113.

Vasanthi, D., and R.K. Mishra, 2008 Epigenetic regulation of genes during development: a conserved theme from flies to mammals. J Genet Genomics 35 (7):413–429.

Wang, W., N. Tindell, S. Yan, and J.H. Yoder, 2013 Homeotic functions of the Teashirt transcription factor during adult Drosophila development. Biol Open 2 (1):18–29.

Weatherbee, S.D., G. Halder, J. Kim, A. Hudson, and S. Carroll, 1998 Ultrabithorax regulates genes at several levels of the wing-patterning hierarchy to shape the development of the *Drosophila* haltere. Genes & Development 12 (10):1474–1482.

Widmann, J., J. Stombaugh, D. McDonald, J. Chocholousova, P. Gardner et al., 2012 RNASTAR: an RNA STructural Alignment Repository that provides insight into the evolution of natural and artificial RNAs. RNA 18 (7):1319–1327.

Wojcik, J., I.G. Boneca, and P. Legrain, 2002 Prediction, assessment and validation of protein interaction maps in bacteria. J Mol Biol 323 (4):763–770.

Zhang, T., F.N. Hsu, X.J. Xie, X. Li, M. Liu et al., 2017a Reversal of hyperactive Wnt signaling-dependent adipocyte defects by peptide boronic acids. Proc Natl Acad Sci U S A 114 (36):E7469–e7478.

Zhang, T., F.N. Hsu, X.J. Xie, X. Li, M. Liu et al., 2017b Reversal of hyperactive Wnt signaling-dependent adipocyte defects by peptide boronic acids. Proceedings of the National Academy of Sciences 114 (36):E7469–E7478.

Zhou, J., H. Ashe, C. Burks, and M. Levine, 1999 Characterization of the transvection mediating region of the *Abdominal-B* locus in *Drosophila*. Development 126 (14):3057–3065.

Zollman, S., D. Godt, G.G. Privé, J.L. Couderc, and F.A. Laski, 1994 The BTB domain, found primarily in zinc finger proteins, defines an evolutionarily conserved family that includes several developmentally regulated genes in Drosophila. Proceedings of the National Academy of Sciences 91 (22):10717–10721.

