## Supplementary material for "Developmental patterning of adipose tissue by *abd-A* and *Abd-B* homeotic genes in *Drosophila melanogaster*": Figures S1-S10

### Supplementary figures

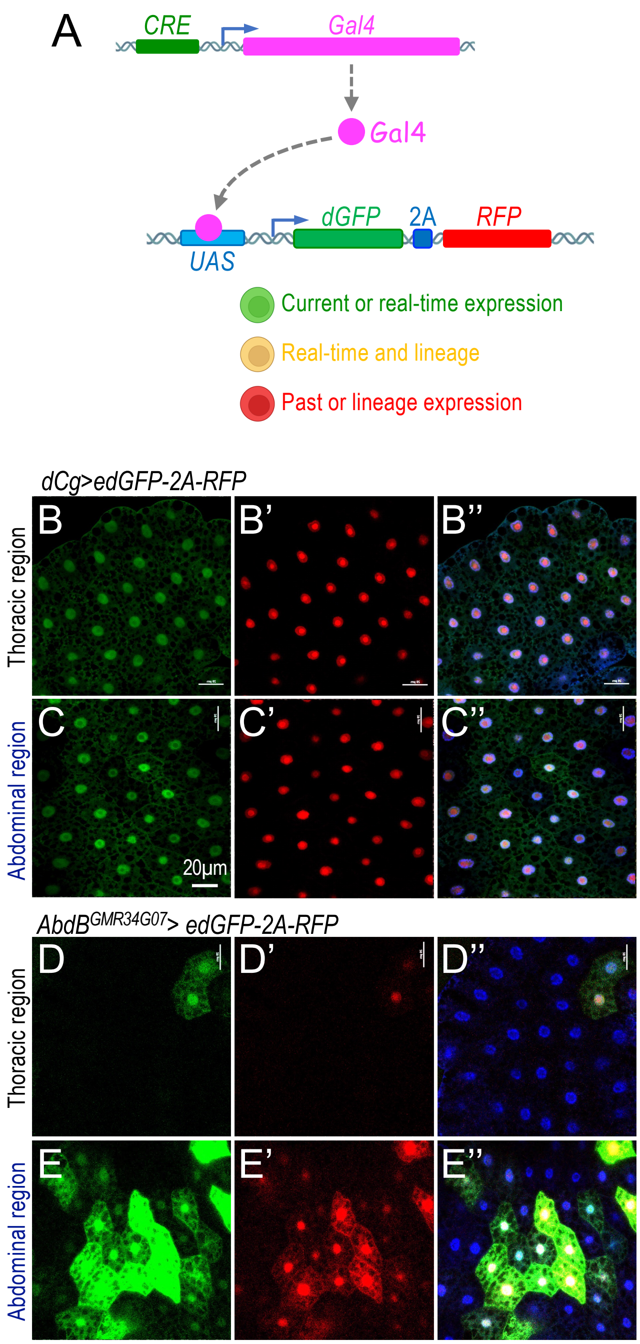


**Fig. S1 Validation of CREs responsible for *Abd-B* expression in the larval fat body.** (A) Schematic diagram illustrating the major steps underlying the UAS-TransTimer system. Upon CRE-driven Gal4 expression, the "TransTimer" cassette, containing destabilized GFP (dGFP) and stable RFP, is activated. In this system, current/real-time expression is marked by green fluorescence, whereas lineage or past expression is indicated by red fluorescence. To further analyze the activity of the *dCg-Gal4* and *AbdB*$^{GMR34G07}$*-Gal4* drivers, we employed the *UAS-TransTimer* reporter system. Genotypes: (B-C'') *dCg-Gal4/+; pUAST-edGFP::2A::RFP/+.* (D-E'') +*; pUAST-edGFP::2A::RFP/AbdB*$^{GMR34G07}$*-Gal4.* (B-B" and D-D") thoracic region; (C-C" and E-E") abdominal region. Green indicates transient or current expression, red indicates lineage or past expression, and blue corresponds to DAPI staining marking nuclei. Scale bar in panel C: 20μm (applies to all images).

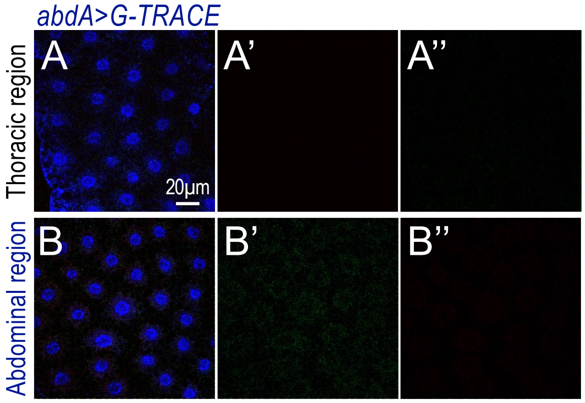


**Fig. S2 Expression pattern of *abdA-Gal4* in the larval fat body.** Lineage tracing analysis of the *abdA-Gal4* driver using the G-TRACE system. Genotype: *+; abdA-Gal4/UAS-RedStinger, UAS-FLP,Ubi-p63E($^{FRT}$STOP$^{FRT}$)GFP.* Red fluorescence represents *RedStinger* expression, indicating current Gal4 activity, whereas green fluorescence indicates GFP expression, marking lineage-tracing cells. (A-A") thoracic region; (B-B") abdominal region. Scale bar in panel A: 20μm (applies to all images).

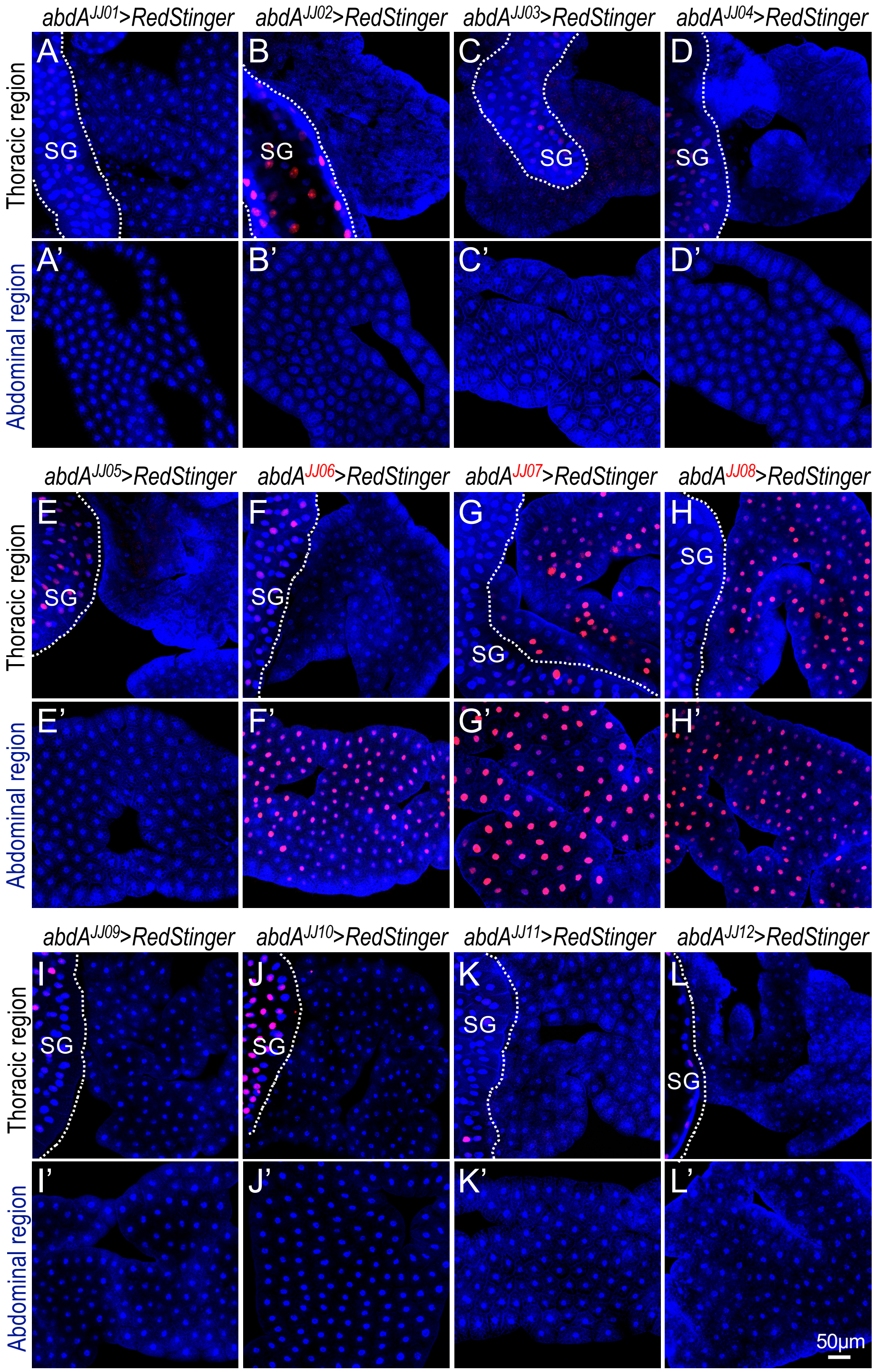


**Fig. S3 Expression patterns of *AbdA-Gal4* drivers generated using the pBPGUw-Gal4 vector.** The *UAS-RedStinger* reporter system was used to examine the expression patterns of the *abdA-Gal4* drivers. Genotypes: (A/A') *UAS-RedStinger/X;* $abdA^{JJ01}$*-Gal4/*+; (B/B') *UAS-RedStinger/X;* $abdA^{JJ02}$*-Gal4/*+; (C/C') *UAS-RedStinger/X;* $abdA^{JJ03}$*-Gal4/*+; (D/D') *UAS-RedStinger/X;* $abdA^{JJ04}$*-Gal4//*+; (E/E') *UAS-RedStinger/X;* $abdA^{JJ05}$*-Gal4/*+; (F/F') *UAS-RedStinger/X;* $abdA^{JJ06}$*-Gal4/*+; (G/G') *UAS-RedStinger/X;* $abdA^{JJ07}$*-Gal4/*+; (H/H') *UAS-RedStinger/X;* $abdA^{JJ08}$*-Gal4/*+; (I/I') *UAS-RedStinger/X;* $abdA^{JJ09}$*-Gal4/*+; (J/J') *UAS-RedStinger/X;* $abdA^{JJ10}$*-Gal4/*+; (K/K') *UAS-RedStinger/X;* $abdA^{JJ11}$*-Gal4/*+; and (L/L') *UAS-RedStinger/X;* $abdA^{JJ12}$*-Gal4/*+*.* The thoracic and abdominal regions are labeled on the left side of the images. SG: Salivary glands. Scale bar in panel A: 50μm (applies to all images).

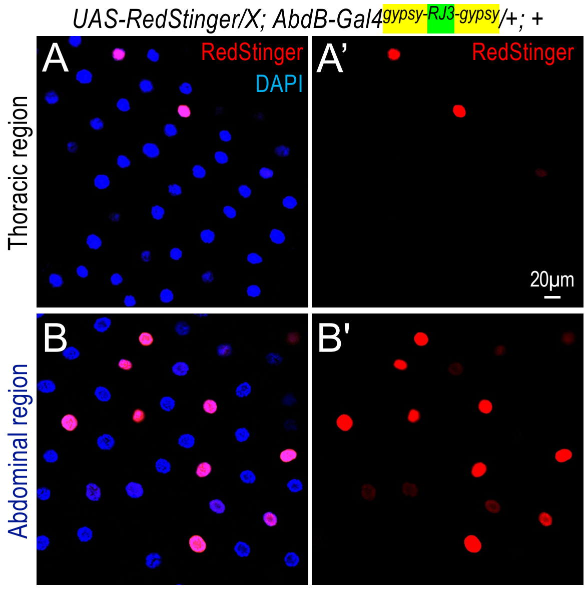


**Fig. S4 The *Abd-B* cis-regulatory element *RJ3* flanked by gypsy insulators exhibits region-specific expression patterns similar to those of *RJ3* without gypsy insulators.** The *UAS-RedStinger* reporter system was used to examine the expression pattern of the *abdB-Gal4* driver containing the RJ3 fragment flanked by gypsy insulators. Genotype: *UAS-RedStinger/X; AbdB-Gal4$^{gypsy\text{-}RJ3\text{-}gypsy}$/+; +*. Red fluorescence indicates RedStinger expression driven by the corresponding *AbdB*-Gal4 construct. Scale bar in panel B: 20µm (applies to all images).

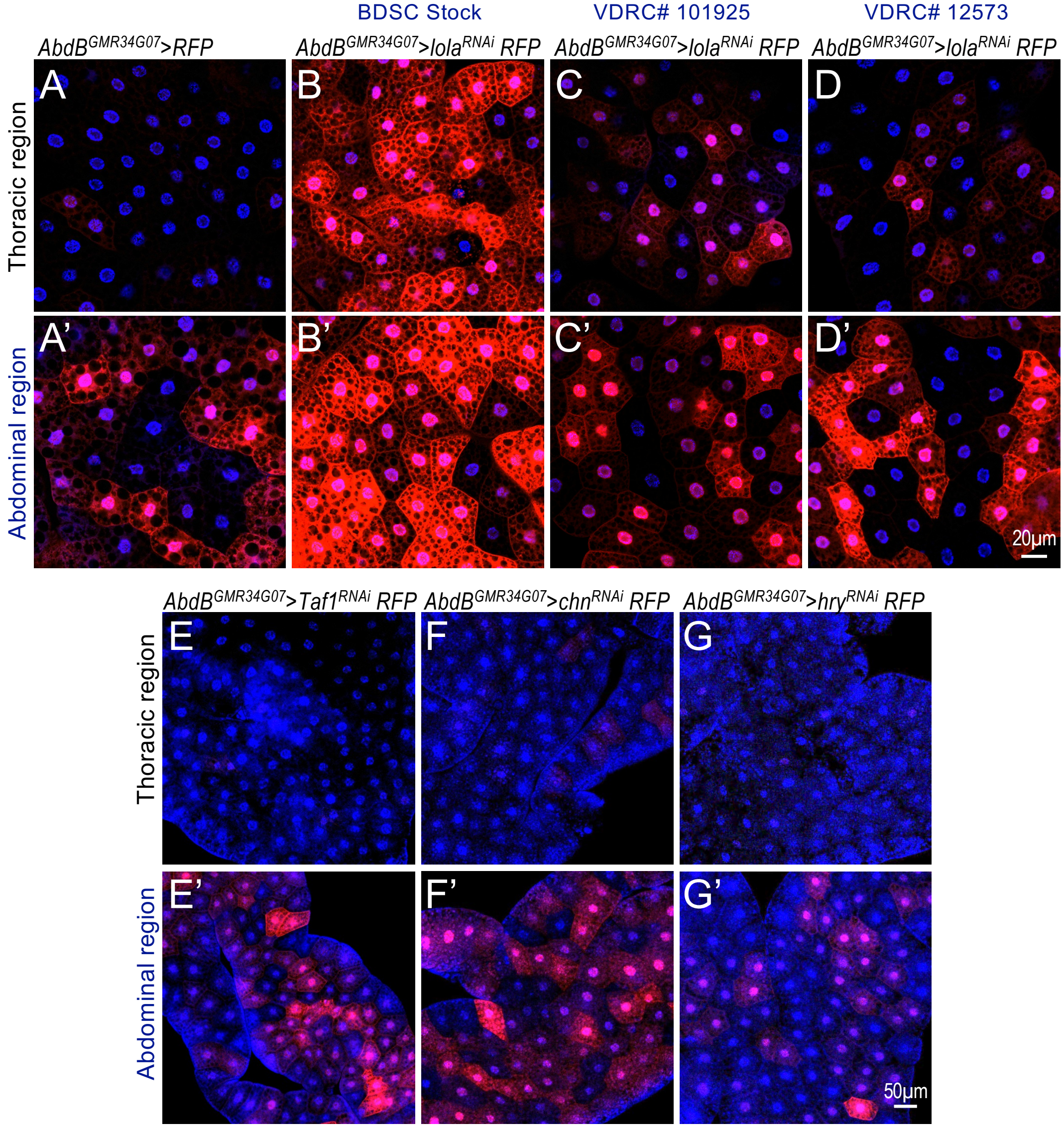


**Fig. S5 Validation of additional Y1H assay candidates using the *UAS-RFP* combined with the *AbdB-Gal4*$^{GMR34G07}$ driver.** (A-D') The effects of depleting *lola* were further validated using two additional RNAi lines from the Vienna *Drosophila* Resource Center (VDRC: #101925 and #12573). The *UAS-RFP/+; AbdB-Gal4*$^{GMR34G07}$*/+* reporter system was used. In all panels, blue indicates DAPI staining marking nuclei, and red indicates RFP expression. Scale bar in panel D': 50μm (applies to all images in A-D'). Genotypes: (A/A') *UAS-RFP/+; AbdB-Gal4*$^{GMR34G07}$/+; (B/B') *UAS-RFP/+; AbdB-Gal4*$^{GMR34G07}$*/UAS-lola*$^{RNAi}$; (C/C') *UAS-RFP/+; AbdB-Gal4*$^{GMR34G07}$*/UAS-lola*$^{RNAi}$ *[VDRC# 101925]* ; *and* (D/D') *UAS-RFP/+; AbdB-Gal4*$^{GMR34G07}$*/UAS-lola*$^{RNAi}$ *[VDRC# 12573].* (E-G') Analysis of *Taf1*, *chn*, and *hry* depletion using the *UAS-RFP* recombined with the *AbdB-Gal4*$^{GMR34G07}$ driver. Genotypes: (E/E') *UAS-RFP/+; AbdB-Gal4*$^{GMR34G07}$*/UAS-Taf1*$^{RNAi}$; (F/F') *UAS-RFP/+; AbdB-Gal4*$^{GMR34G07}$*/UAS-Chn*$^{RNAi}$; and (G/G') *UAS-RFP/+; AbdB-Gal4*$^{GMR34G07}$*/UAS-Hry*$^{RNAi}$. The thoracic and abdominal regions are labeled on the left side of the images. Scale bar in panel G': 50 μm (applies to images E-G').

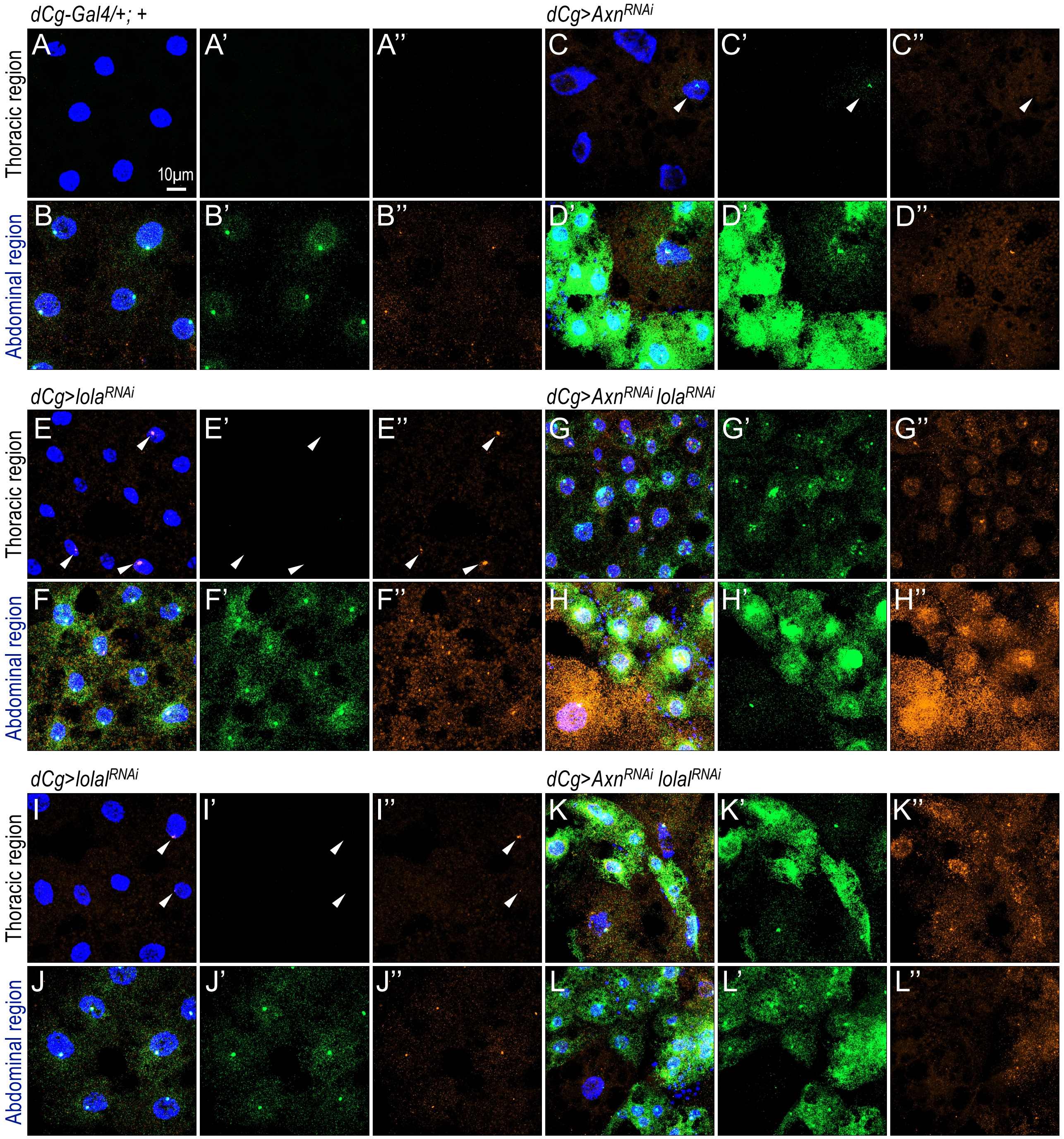


**Fig. S6: Analysis of *abd-A* and *Abd-B* transcript levels following depletion of *lola* or *lolal* under conditions of activated Wnt signaling.** HCR RNA-FISH was used to detect *abd-A* (green) and *Abd-B* (orange) mRNA transcripts in larval adipocytes. White arrowheads indicate thoracic adipocytes expressing *abd-A* or *Abd-B* transcripts. Genotypes: (A-A'') and (B-B'') *dCg-Gal4/+;* +; (C-C'') and (D-D'') *dCg-Gal4/UAS-Axn*$^{RNAi}$*;* +; (E-E'') and (F-F'') *dCg-Gal4/+; UAS-lola*$^{RNAi}$/+; (G-G'') and (H-H'') *dCg-Gal4/UAS-Axn*$^{RNAi}$*; UAS-lola*$^{RNAi}$/+; (I-I') and (J-J'') *dCg-Gal4/+; UAS-lolal*$^{RNAi}$/+; and (K-K'') and (L-L'') *dCg-Gal4/ UAS-Axn*$^{RNAi}$*; UAS-lolal*$^{RNAi}$*/+.* The thoracic and abdominal regions are labeled on the left side of the images. Scale bar in panel A: 10μm (applies to all images).

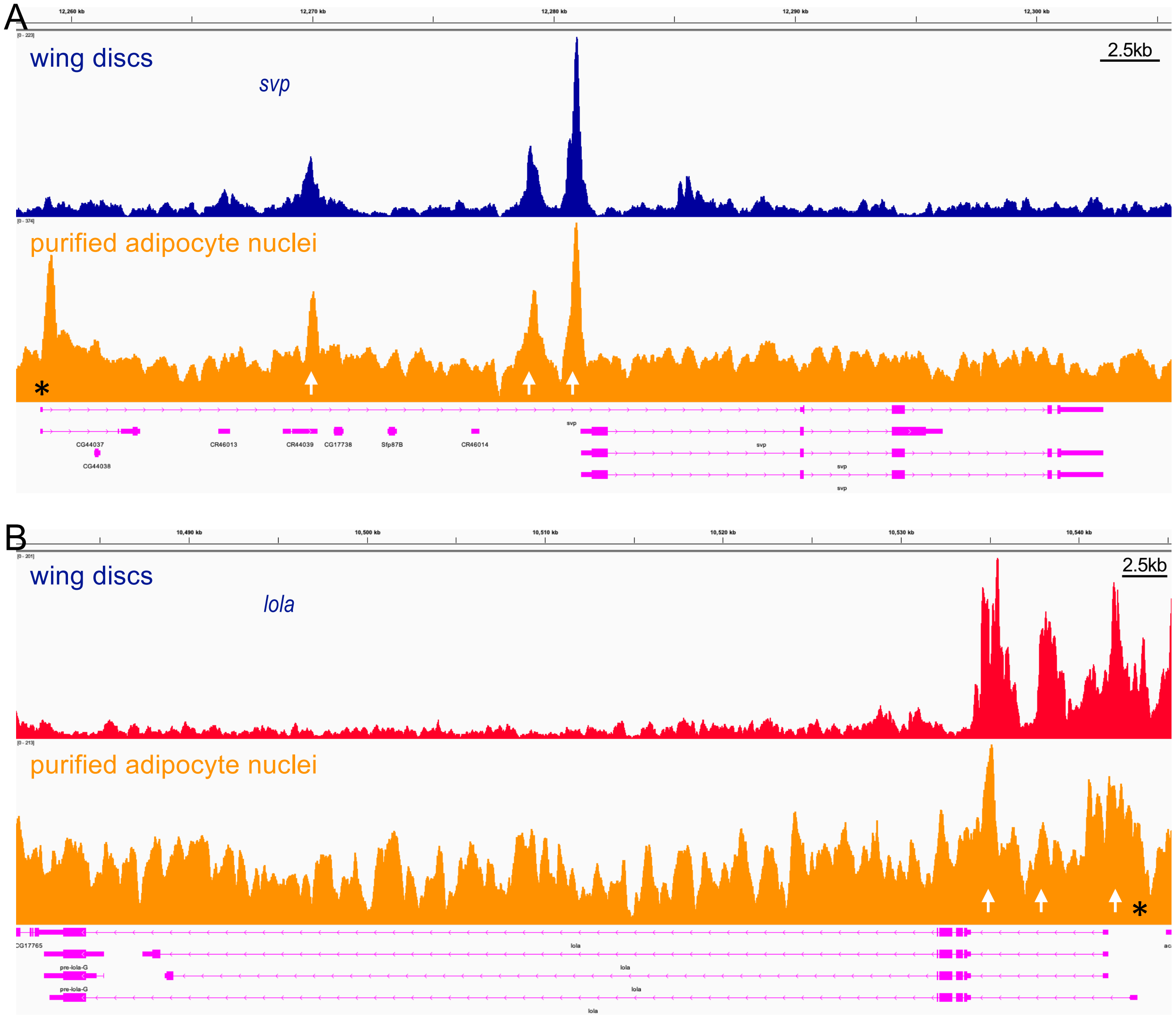


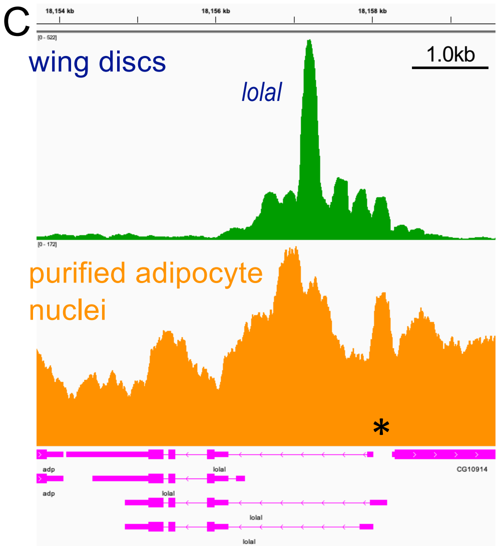


**Fig. S7: Genome browser tracks displaying dTCF/Pan binding peaks at the *svp* (A), *lola* (B), and *lolal* (C) loci.** Peaks were identified by CUT&RUN analysis performed on wing imaginal discs (upper panels, dark blue indicating downregulated expression and red indicating upregulated expression upon Wnt/Wg activation) and purified adipocyte nuclei from larval fat bodies (lower panels, orange). Tracks are visualized in the IGV_2.14.1 browser. The y-axis is autoscaled, and four gene isoforms per locus are displayed in magenta. Shared prominent peaks between datasets are indicated by arrows. (*) indicates transcription start sites.

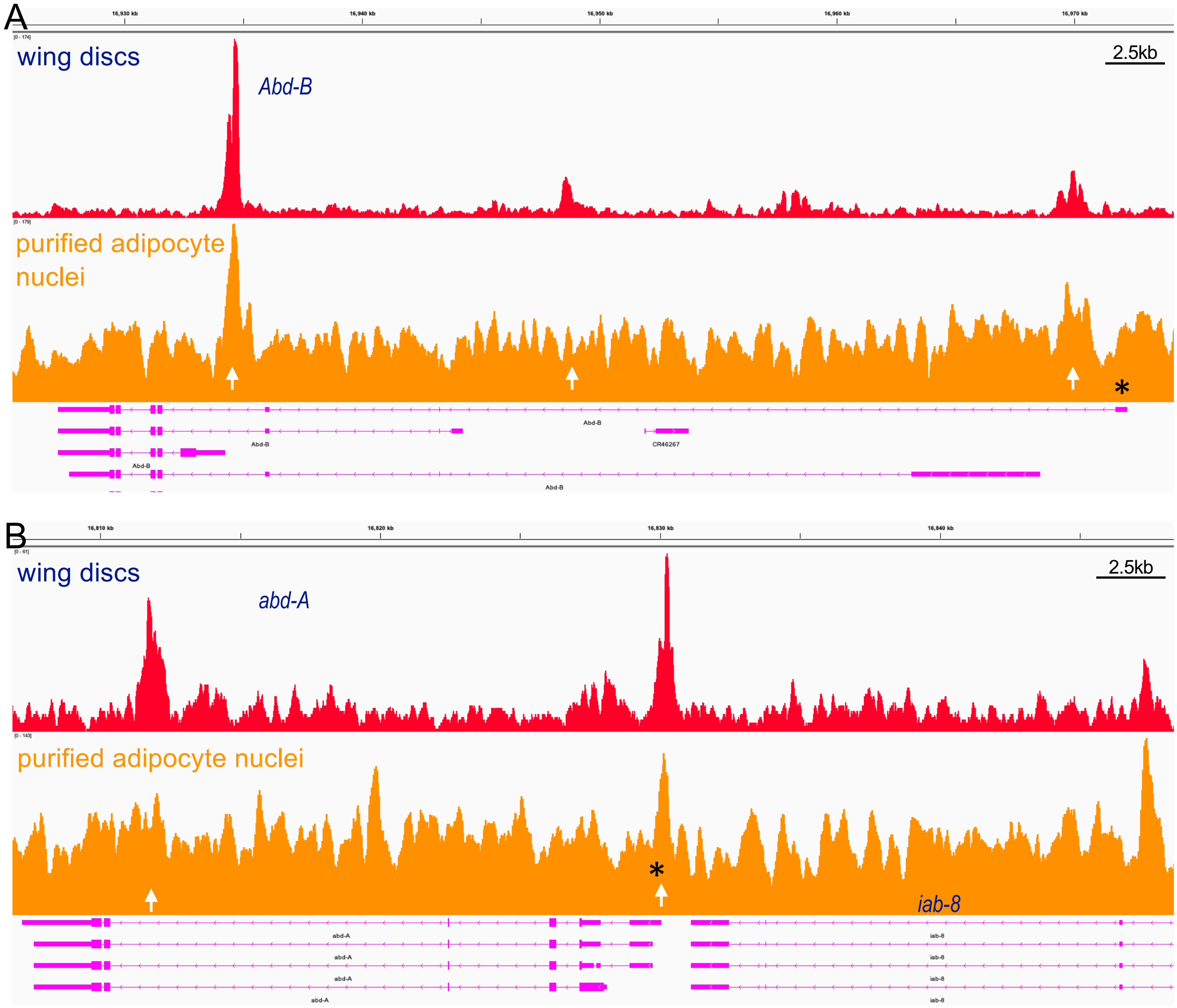


**Fig. S8: Genome browser tracks showing dTCF/Pan binding peaks at the *Abd-B* (A) and *abd-A* (B) loci.** Peaks were identified by CUT&RUN analysis performed on wing imaginal discs (upper panels, red indicating upregulated expression upon Wnt/Wg activation) and purified adipocyte nuclei from larval fat bodies (lower panels, orange). Tracks are visualized in the IGV_2.14.1 browser. The y-axis is autoscaled, and four gene isoforms per locus are displayed in magenta. Shared prominent peaks between datasets are indicated by arrows. (*) indicates transcription start sites. Scale bars: 2.5 kb.

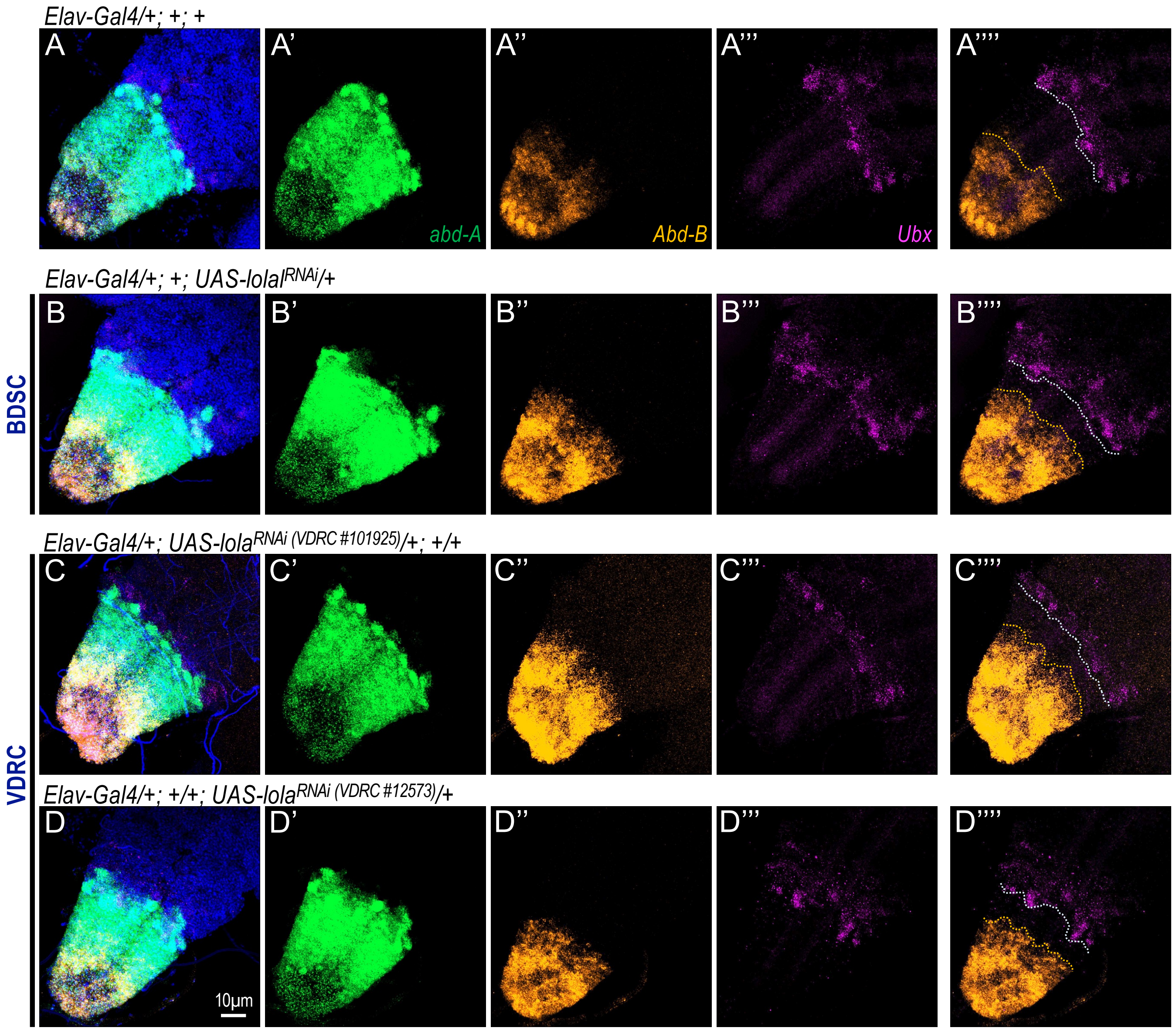


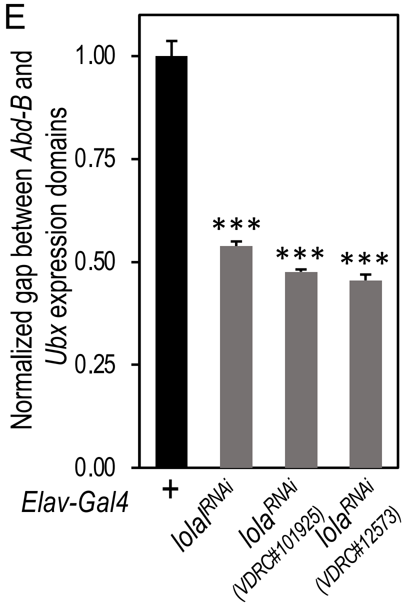


**Fig. S9: *Elav-Gal4*-driven depletion of *lola* or *lolal* markedly expands the *Abd-B* expression domain in the larval ventral nerve cord (VNC).** HCR RNA-FISH was used to detect *abd-A* (green), *Abd-B* (orange), and *Ubx* (magenta) mRNA transcripts in the larval VNC. Scale bar in panel D: 10μm (applies to all images). Genotypes: (A-A””) *UAS-RFP/+; AbdB-Gal4*$^{GMR34G07}$*/+*; (B-B””) *UAS-RFP/+; AbdB-Gal4*$^{GMR34G07}$*/UAS-lolal*$^{RNAi}$; (C-C””) *UAS-RFP/+; AbdB-Gal4*$^{GMR34G07}$*/UAS-lola*$^{RNAi}$ *[VDRC #101925]*; and (D-D””) *UAS-RFP/+; AbdB-Gal4*$^{GMR34G07}$*/UAS-lola*$^{RNAi}$ *[VDRC #12573].* (E) Quantification of the normalized gap between *Abd-B* and *Ubx* expression domains in the VNC (n = 3 independent biological replicates, with 5 independent measurements obtained from each VNC). P values were calculated using one-tailed unpaired *t*-tests, based on predefined directional predictions in the experimental design. Error bars represent standard deviations. ***: P<0.001.

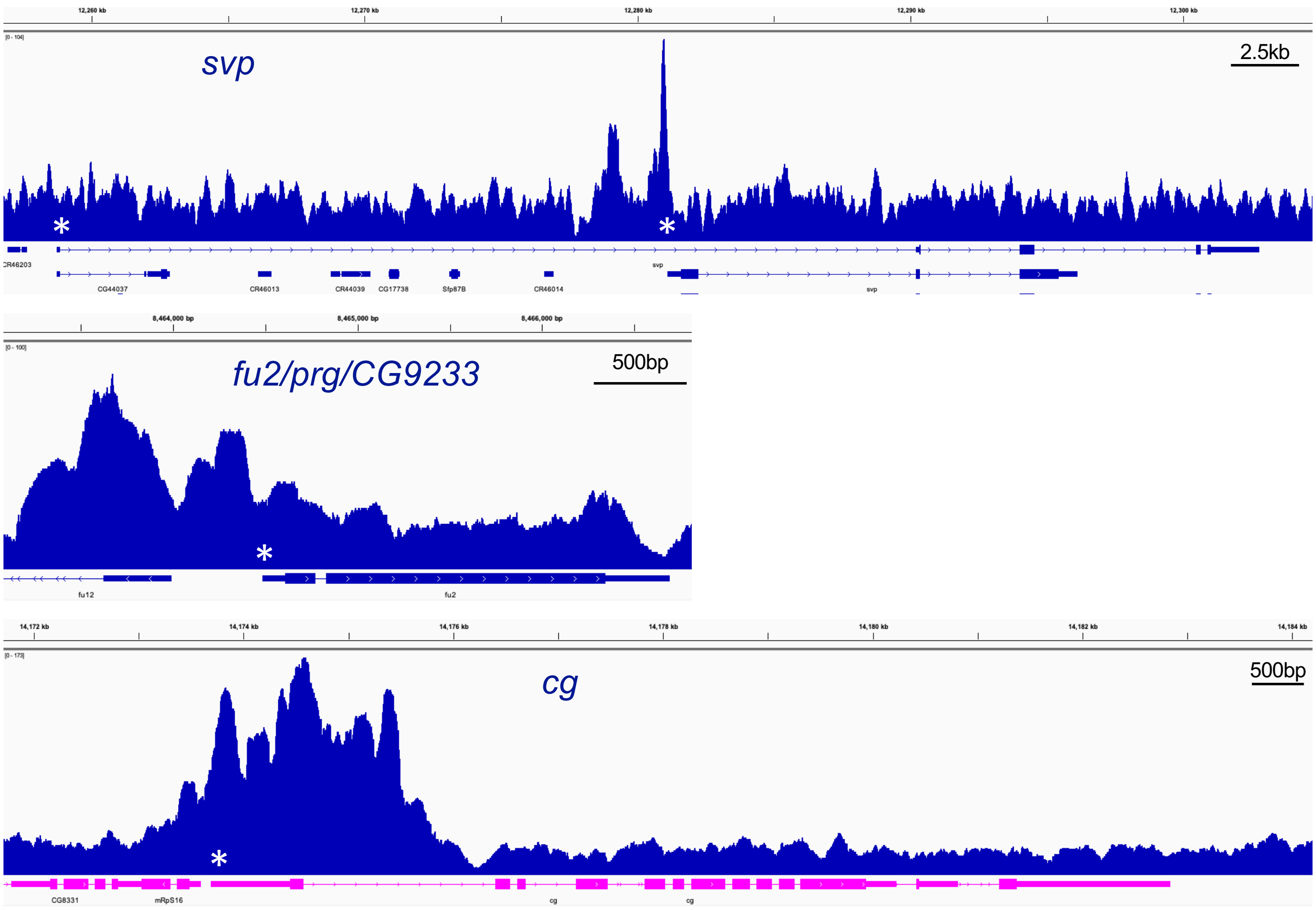


**Fig. S10: Genome browser tracks showing Abd-B binding peaks at the *svp* (A), *prg* (B), and *cg* (C) loci.** Peaks were identified by CUT&RUN analysis performed on larval central nervous system (CNS), with enrichment primarily in the ventral nerve cord (VNC). Tracks are visualized in the IGV_2.14.1 browser. The y-axis is autoscaled. (*) indicates transcription start sites.

### Supplementary Tables

**Table S1.** Table S1 Gal4 drivers within the Abd-B locus and their activities in the larval fat body.

**Table S2.** Newly generated Gal4 drivers within the *abd-A* locus and their activities in adipocytes.

**Table S3.** *Drosophila* stocks used in this study.

**Table S4.** Primers used in this study.
